# Evidence that the cell glycocalyx envelops respiratory syncytial virus (RSV) particles that form on the surface of RSV-infected human airway cells

**DOI:** 10.1101/2024.10.08.616916

**Authors:** Soak Kuan Lai, Zhi Qi Lee, Trina Isabel Tan, Boon Huan Tan, Richard J Sugrue

**Author notes:** Corresponding author; /.

## Abstract

We examined how respiratory syncytial virus (RSV) particles circumvent the overlying glycocalyx on virus-infected A549 cells. The glycocalyx was detected using the lectin WGA-AL488 probe, and the antibodies anti-HS and anti-syndecan-4 that detect heparin sulphate (HS) and the syndecan-4 protein (SYND4) respectively. Imaging of RSV-infected cells provided evidence that the glycocalyx envelopes the virus filaments as they form, and that components of the glycocalyx such as HS moieties and SYND4 are displayed on the surface of the mature virus filaments. Using recombinant expression of the G protein we also demonstrated that the G protein was trafficked into pre-existing filamentous cellular structures with a well-defined glycocalyx, suggesting that the glycocalyx is maintained at the site of virus particle assembly. These data provide evidence that during RSV particle assembly the virus filaments become enveloped by the glycocalyx, and that the glycocalyx should be considered as a structural component of RSV particles.

## Introduction

Human respiratory syncytial virus (RSV) is currently the most important viral cause of lower respiratory tract infection in young children and neonates, leading to high levels of mortality and morbidity [1]. On the surface of RSV-infected cells the mature RSV virus particle assembles as cell-associated filamentous structures[2], which we refer to as virus filaments. The virus filaments are bounded by a lipid envelope that is derived from the host cell into which the virus-encoded G and F glycoproteins are inserted, and beneath the virus envelope is a protein layer formed by the Matrix (M) protein. While the G protein is established as the RSV cell-attachment protein [3, 4], the F protein initiates fusion of the virus envelope and cell membrane during virus entry into the host cell. The F protein is initially expressed as a single polypeptide chain (F0), which is subsequently cleaved by furin into the F1 and F2 subunits [5–7] prior to being inserted into the virus envelope. The interaction between the G and F protein has been demonstrated on virus particles[8, 9], and it is expected that their concerted biological activities are required for mediating efficient virus transmission. The virus genomic RNA (vRNA) is packaged in the virus filaments as part of a larger structure called the virus nucleocapsid (NC). The virus nucleocapsid (N) protein coats the vRNA in the NC and it is the most abundant virus protein present in the NC, but the biologically active form of the NC also requires the presence of the virus phosphoprotein (P protein), the M2-1 protein and the large (L) protein [10–13]. Virus filament formation occurs in many permissive cell lines, and that they form on organoid airway cell systems that are infected with RSV[14] suggests that virus filament formation has a clinical relevance. In these cell systems the virus filaments mediate virus transmission [9, 14–18], and experimental conditions that impair virus filament formation also inhibit RSV transmission [15–17, 19].

Interactions between the different virus structural proteins have been demonstrated, and it is presumed that multi-protein complexes involving these virus proteins facilitate the stabilization of the mature virus particles. However, several host cell factors have also been shown to play important roles in virus filament formation. Specialised lipid domains called lipid raft microdomains and F-actin structures are present at the site of virus particle assembly, and the available evidence suggests the involvement of specialised lipid raft microdomains called caveolae in this process[16, 19–24]. Although the lipid raft microdomains may impart important biophysical properties to the viral envelope, raft-associated host cell factors also play a direct role in virus particle assembly. Virus-induced activation signalling pathways that are raft-associated also modulate F-actin structures, and collectively these cellular factors have been shown to play a role in the process of RSV particle morphogenesis[15, 19, 25, 26]. These previous studies highlight the important role that cellular factors and process play in the formation and the morphogenesis of the mature and infectious RSV particles.

Although the role that the specialised domains within the plasma membrane play in virus assembly has been demonstrated, a feature of the host cell interaction on the cell surface that has not been fully examined during RSV particle assembly is the glycocalyx. The glycocalyx is a complex polysaccharide-rich matrix layer that forms a relatively thin viscous coating on the surface of different tissues and cell types [27–29]. The glycocalyx extends from the cell surface into overlying extracellular matrix, and depending on the nature of the cell type in question, the thickness of the glycocalyx can range from between 1nm to up to several hundred nm’s in thickness [30, 31].. An array of biological molecules form the glycocalyx, and these arise as the result of different cellular processes that occur in the cell, giving rise to a complex structure that is often difficult to define biochemically [30–34]. The glycocalyx consists of several different biomolecules and include high levels of glycoaminoglycans such as heparan sulphate and hyaluronic acid. In addition, specific proteoglycans are also present in the glycocalyx, including the glycosylphosphatidylinositol (GPI) anchored protein glypican and the transmembrane protein syndecan-4 (SYND-4) whose ectodomains are post-translationally modified by the attachment of heparan sulphate (HS) molecules [35, 36]. While the glycocalyx was traditionally thought to function by providing a protective covering on the surface of cells[37], the increasing importance of the glycocalyx in an array of other physiological processes on cells and tissues is being becoming realised. For example, recent evidence suggests that the glycocalyx also plays a role in controlling cell morphology and in regulating the diffusion of proteins in the cell membrane[31, 38, 39]. Evidence suggests that the cell glycocalyx can be disrupted in cells infected by several different viruses and that this change in the glycocalyx may be associated with severity of infection [40–42]. The glycocalyx is present on airway tissue and on the epithelial cells that are permissive for RSV infection, however it is currently unclear how the RSV particles circumvent this potential cell barrier during the process of virus particle assembly and egress. HS has been proposed to play a role in facilitating the attachment of the virus to the host cell during the initial stages of the RSV replication cycle [43, 44]. Since HS is a major component of the glycocalyx, this suggests that the glycocalyx may also play a role in the RSV replication cycle. There are inherent difficulties in isolating and biochemically analysing components of the cell glycocalyx, therefore in this current study we have employed *in situ* imaging to examine how the newly assembled RSV particles virus circumvent the overlying glycocalyx during RSV particle assembly.

## MATERIALS AND METHODS

### Virus and cell preparation

The RSV A2 virus was prepared in HEp-2 cells as described previously [45]. The HEp2 and A549 cells were maintained at 37°C in Dulbecco’s Modified Eagle Medium (DMEM) supplemented with 10% FCS in a humidified chamber with 5% CO_2_.

### RSV infection

This was performed as described previously [18]. Briefly, cells that were 95% confluent were infected with RSV using the required multiplicity of infection. Infections were performed in cells using DMEM supplemented with 2% FCS in a humidified chamber at 33 °C with 5% CO_2._ At specific time intervals the cells were harvested and processed further.

### Antibodies and specific reagents

The anti-mouse and anti-rabbit IgG conjugated to Alexa Fluor™ 488 and Alexa Fluor™ 555 (ThermoFisher), the mouse monoclonal antibody anti-HS/clone F58-10E4 (Amsbio), the rabbit polyclonal antibody anti-syndecan-4/ab74139 (abcam), and wheat germ agglutinin conjugated to Alexa Fluor™ 488 (ThermoFisher) were purchased. The RSV N and M protein antibodies have been described previously [20, 46, 47], and the F protein antibody and G protein antibody were gifts from Jose Melero (Madrid, Spain) and Geraldine Taylor (IAH, UK) respectively.

### Recombinant virus protein expression

The construction of the G, F and M protein expression vectors has been described previously. Briefly, the G, F and M gene of the RSV A2 strain was cloned into the recombinant pCAGGS vector to create pCAGGS/G, pCAGGS/F and pCAGGS/M respectively. Bulk preparation of expression vectors was performed using the plasmid Midiprep kit (Qiagen). Cells (1LJ×LJ10^5^ cells per well) in a 24-well cluster plate were transfected with 1 μg plasmid DNA using the Lipofectamine 3000 system (Invitrogen) following the manufacturer’s instructions. The transfected cells were incubated at 33°C in a humidified chamber with 5% CO_2_ and at 20 hrs post-transfection the cells were processed further.

### Western blotting

The transfected cells were washed using ice cold PBS (4 °C) and extracted directly into Boiling Mix (1% SDS, 5% mercaptoethanol in 20 mM Tris/HCL, pH 7.5). After heating at 95°C for 2 min the cell extracts were clarified by centrifugation (13,000×g for 2 min) and the proteins separated by SDS-PAGE and transferred by Western blotting onto nitrocellulose membranes. In all cases the apparent molecular masses were estimated using Kaleidoscope protein standards (Bio Rad, USA). The protein bands were visualized using the ECL protein detection system (Amersham, UK).

### Immunofluorescence microscopy

The cells on 12-mm circular glass coverslips were fixed with 4% paraformaldehyde (Sigma-Aldrich) and washed with PBS. The cells were either non-permeabilized and antibody stained or permeabilized using 0.1% Triton X-100 in PBS at 4°C for 15 min prior to antibody staining. The cells were stained with the appropriate primary and secondary antibody combinations and mounted on microscope slides using CitiFluor. Surface labelling using Wheat germ agglutinin conjugated to Alexa Fluor™ 488 (WGA-AL488) was performed according to the manufacturer’s instructions. For immunofluorescence microscopy imaging, the stained cells were imaged using a Nikon Eclipse 80i Microscope (Nikon Corporation, Tokyo, Japan) with an Etiga 2000R camera (Q Imaging, Teledyne Photometrics, Tucson AZ, USA) attached. The images of immunofluorescence-stained cells were recorded using Q Capture Pro ver. 5.0.1.26 (Q Imaging, Teledyne Photometrics). Imaging for confocal microscopy was performed using a Zeiss 910 confocal microscope (Zeiss, Oberkochen, Germany) with Airyscan 2 processing using the appropriate machine settings. In all cases, a series of consecutive images were obtained in the Z projection (Z-stack) from the stained cells in the field of view. The recorded images were examined and processed using Zen ver. 2.3 software (Zeiss).

## Results and Discussion

### 1. Detecting the glycocalyx on the surface of A549 cells

In the current study we primarily used A549 cells, which is an established human airway cell line used to examine RSV infection. Three established reagents were used to detect the glycocalyx on the A549 cells, the probe Wheat germ agglutinin conjugated to Alexa Fluor™ 488 (WGA-AL488), anti-HS antibody that detects the presence of Heparin Sulphate (HS) and the anti-SYND4 antibody which detects the presence of the syndecan-4 (SYND4) protein. The WGA-AL488 is an established cellular probe that is used to image the surface of cells. The WGA-AL488 is a lectin that binds sugars (N-acetyl-D-glucosamine and sialic acid), and it is therefore able to delineate the cell surface by labelling the carbohydrate components of the glycocalyx that form on the cell surface. Therefore, it is also an established cellular probe that can be used to visualise the glycocalyx in different tissues and cell types (e.g. [48, 49]) Although the WGA-AL488 probe will label specific glycan components of the glycocalyx rather than all the different sugar moieties that are present in the glycocalyx, the WGA-AL488 probe nevertheless allows detection of the glycocalyx. This probe has recently been used to examine the effect of drug treatment on the glycocalyx in A549 cells [50]. However, the glycocalyx is a complex structure that contains several different biological molecules, and the presence of high levels of heparan sulphate (HS) and the SYND4 protein in the glycocalyx is established. The antibody anti-HS (F58-10E4) was prepared using HS derived from lung tissue and it is an established immunological reagent that is used to detect HS in biological tissues and cells [51–55], and we used anti-HS to detect the HS on A549 cells,. The HS chain most commonly consists of repeating disaccharide unit composed of a glucuronic acid (GlcA) linked to N-acetylglucosamine (GlcNAc), and the recognition by the anti-HS monoclonal antibody requires the N-sulfation of GlcNAc residues at the antibody binding epitope in the HS chain (https://resources.amsbio.com/Guide/Heparan-Sulfate-Antibodies-Application-Guide.pdf).

The protein SYND4 is a HS-modified type 1 transmembrane protein found on the surface of many epithelial cell types, and it is an established component of the glycocalyx. The antibody anti-SYND4 is an established immunological reagent used to detect the SYND4 protein e.g. [56], and in this study it was used to detect the presence of SYND4 protein on the surface of A549 cells.

We initially used these reagents to examine the glycocalyx on mock-infected A549 cells i.e. prior to virus infection. Non-permeabilised mock-infected cells were stained with WGA-AL488 and examined using immunofluorescence (IF) microscopy to visualise the distribution of the glycocalyx on the surface of the A549 cells (Fig. 1A). This showed widespread WGA-AL488 staining across the entire cell monolayer, consistent with the glycocalyx extending across the whole cell monolayer. A more detailed analysis of the WGA- AL488-stained non-permeabilised mock-infected cells was performed by confocal microscopy that allowed detailed imaging of the surface of representative A549 cells (Fig. 1B (i) and C(i)). A detailed analysis indicated low levels of WGA-AL488 staining on the flat regions of the cell surface together with higher levels of a filamentous WGA-AL488 staining (Fig. 1B(ii) and C(ii)). The latter staining pattern was consistent with the labelling of surface projections such as the microvilli that form on the surface of these cells. The series of images from the same WGA-AL488-stained non-permeabilised infected cell that was captured in the Z-projection was reconstructed into single 3-dimensional images (Fig. 1D (i) and (ii)), which further highlighted the WGA-AL488 staining on the cell surface projections on these cells.

**Figure 1.**
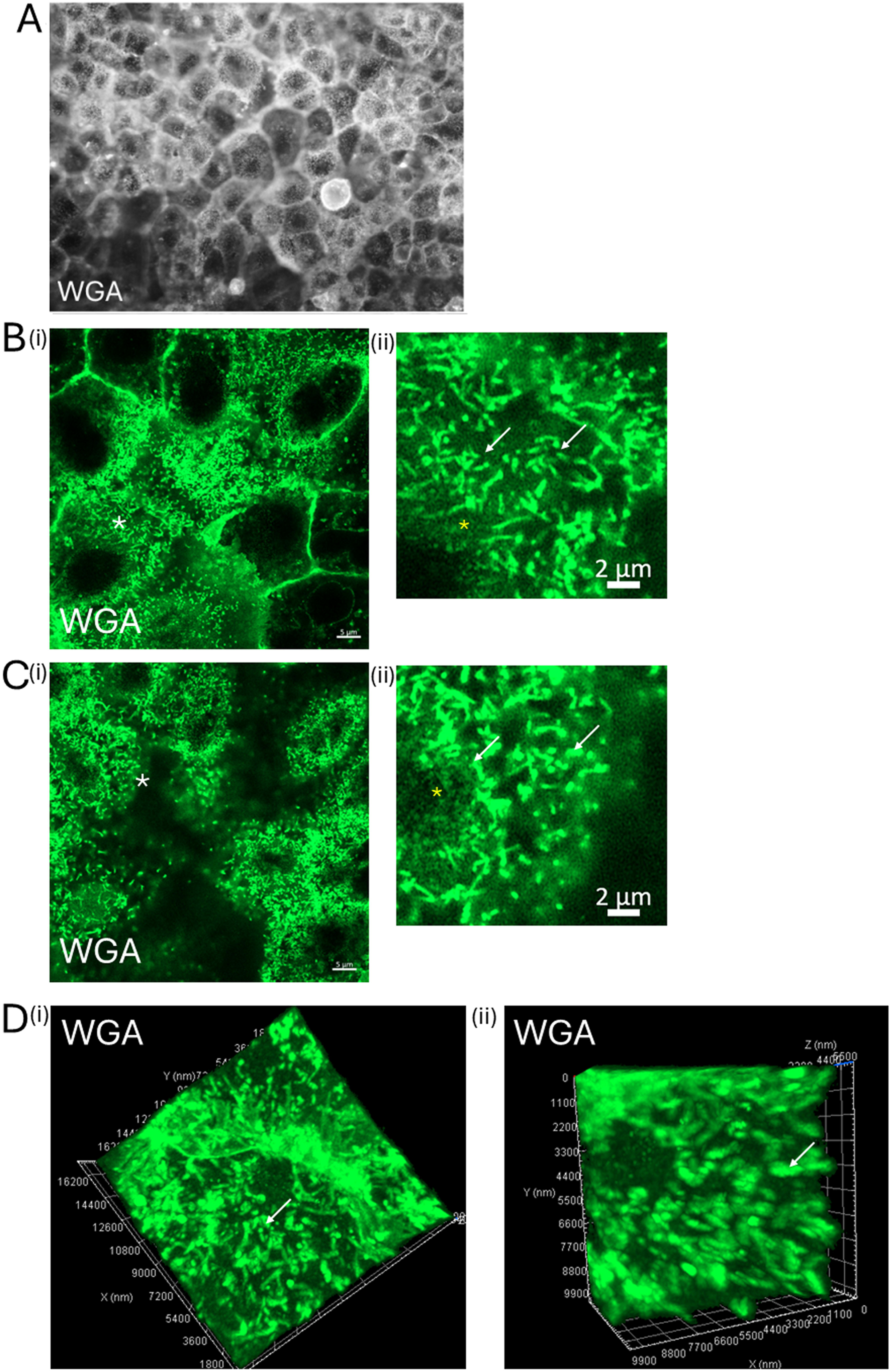
Distribution WGA-AL488-stained glycocalyx on the surface of mock-infected A549 cells. A549 cell monolayers were stained using WGA-AL488 (WGA) and **(A)** imaged using immunofluorescence microscopy (magnification x40 objective). **(B and C)** Representative WGA-AL488 stained cells were imaged using confocal microscopy and a series of images in the Z-plane (Z-stack) were recorded. Images at **(B)** (i) and (ii) the cell periphery and **(C)** (i) and (ii) close to the cell top from the image series are shown. In each case (ii) are enlarged images taken from the area highlighted (white *) in (i). **(D)** (i) and (ii) A series of images through the WGA-AL488-stained cell that was obtained in the Z projection were processed into a single 3-dimentional images (ii) is an enlarge image taken from (i). The WGA-AL488-stained surface projection (white arrows) and flat regions (yellow *) are highlighted.

Although staining was observed on the flat areas on the cell surface (devoid of filamentous projections), the WGA-AL488 stained the surface of cellular projections such as microvilli suggesting that the glycocalyx was localised on these surface projections. The glycocalyx can generate mechanical forces on the cells plasma membrane that determine the shape of cells, and in this context the glycocalyx can regulate the formation of actin-stabilized membrane projections of the surface of cells such as microvilli and filopodia [39].

The WGA-AL488 staining pattern that we observe on A549 cells is consistent with recent studies using chemical labelling procedure have demonstrated the presence of the glycocalyx on microvilli and filopodia [30].

Non-permeabilised mock-infected cells were also co-stained using WGA-AL488 and anti-HS and examined by IF microscopy. This allowed the respective distribution of the WGA-AL488 and anti-HS staining to be imaged, which showed widespread WGA-AL488 and anti-HS staining across the surface of the entire cell monolayer (Fig. 2A). Representative WGA-AL488 and anti-HS co-stained cells were examined in more detail using confocal microscopy that allowed imaging of the cell periphery (Fig. 2B(i)) and cell top (Fig. 2C(i)), which confirmed anti-HS staining on the surface of the WGA-AL488-stained cells. This analysis confirmed both the presence of anti-HS staining on the WGA-AL488-stained filamentous projections at both the cell periphery (Fig. 2B(ii)) and cell top (Fig. 2C(ii)), as well as the anti-HS staining on the flat regions of the cell membrane. The series of images in the Z-projection from the same WGA-AL488 and anti-HS co-stained non-permeabilised infected cell was reconstructed into single 3-dimensional images (Fig. 2D). This further highlighted the anti-HS staining on the WGA-AL488 stained on the cell surface projections, as well as the presence of the anti-HS staining on the flat regions of the cell membrane. The non-permeabilised A549 cells were also co-stained with anti-HS and anti-SYND-4 and examined by confocal microscopy (Fig 2D). The anti-SYND-4 exhibited a diffuse staining pattern across the cell surface that was similar in appearance to the staining pattern exhibited by anti-HS and confirmed that the SYND4 protein was expressed on the surface of these cells.

**Figure 2.**
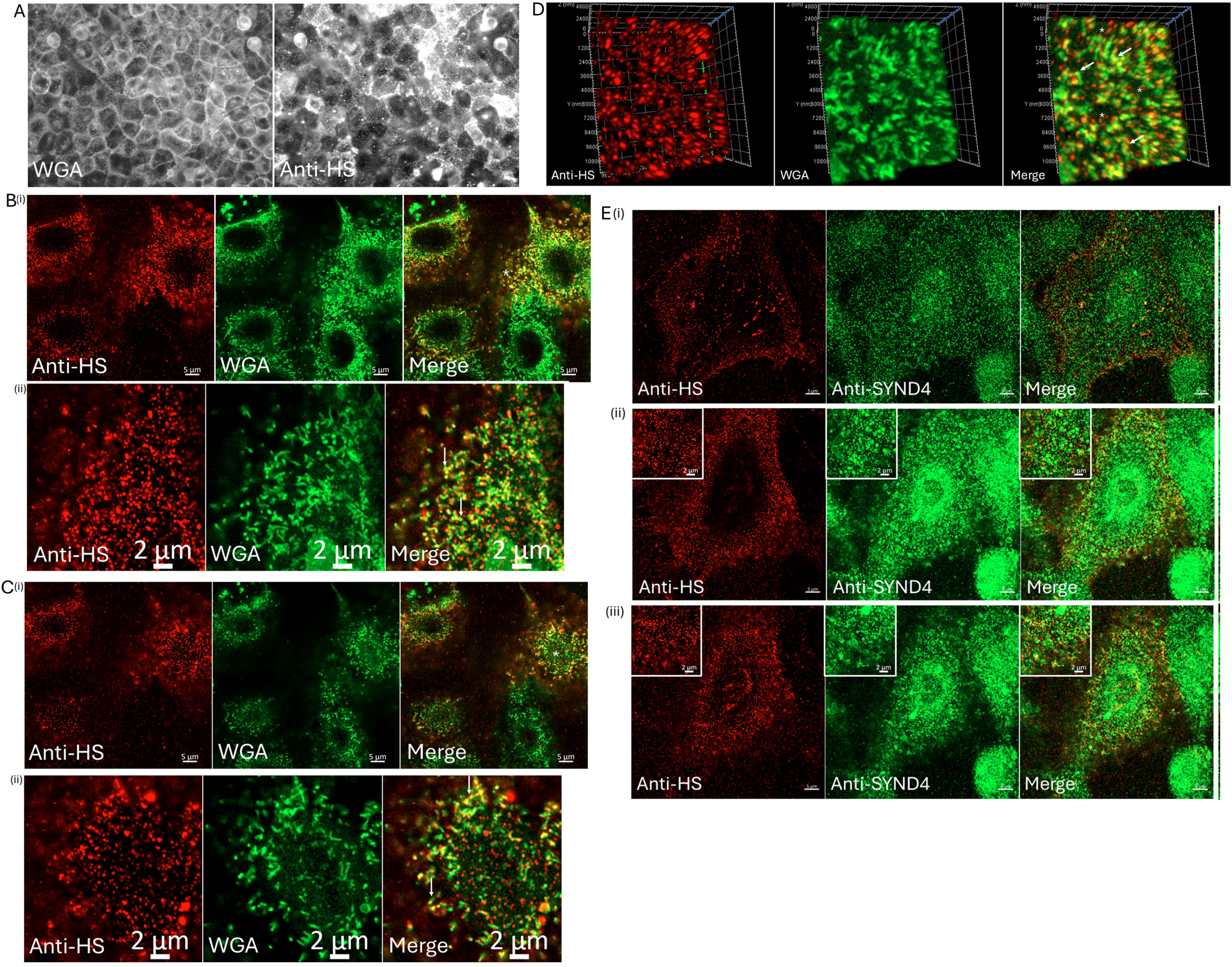
Distribution of the heparan sulphate and Syndecan-4 (SYND4) on the surface of mock-infected A549 cells. A549 cell monolayers were co-stained using WGA-AL488 (WGA) and anti-heparan sulphate antibody (anti-HS). The co-stained cells were imaged using **(A)** immunofluorescence microscopy (magnification x20 objective) and **(B and C)** representative WGA-AL488 and anti-HS-stained cells were imaged using confocal microscopy and a series of images in the Z-plane (Z-stack) were recorded. Images at **(B)** (i) and (ii) the cell periphery and **(C)** (i) and (ii) close to the cell top from the image series are shown. **(B)** (ii) and **(C)** (ii) are enlarged images taken from the area highlighted (white *) in **(B)** (i) and **(C)** (i) respectively. The WGA-AL488 and anti-HS co-stained surface projection (white arrows) are highlighted. **(D)** A series of images through the WGA-AL488 and anti-HS co-stained cells that was obtained in the Z projection were processed into a single 3-dimentional image. The anti-HS staining on the filamentous projections (white arrows) and on the flat regions of the cell brane (*) are highlighted. **(E)** A549 cell monolayers were co-stained using anti-SYND4 and anti-HS and the co-stained cells imaged using confocal microscopy and a series of images in the Z-plane (Z-stack) were recorded. Representative anti-SYND4 and anti-HS-stained cells are presented that were imaged at (i) the cell periphery and (ii) close to the cell top and (iii) at the cell top. Insets are enlarged images taken from the main plate. In the confocal microscopy images the individual image channels and the merged images are shown.

Collectively, the data that was obtained using these different probes was consistent with the presence of a developed glycocalyx on the surface of A549 cells prior to infection. The glycocalyx was located on both clearly visible cellular projections e.g. microvilli and on the flat regions of the cell surface. Interestingly, in general the WGA-AL488-staining was more intense on the filamentous projections than on the flat regions of the cell, suggesting that there way be differences in the glycocalyx composition at these different locations.

### 2. The anti-F staining on the surface of RSV-infected cells delineates the filamentous virus particles (virus filaments)

In this study the A549 cells were infected with RSV using a multiplicity of infection (moi) of 0.05, and unless otherwise indicated, the infected cells were examined at 24 hrs post-infection (hpi). This is at a sufficient time of infection for the virus filaments to form on the surface of RSV-infected A549 cells using these experimental conditions[9, 18]. Prior to examining the relationship between the glycocalyx and virus particles, we first confirmed that staining with antibodies that recognise the RSV F protein could be used to detect the RSV particles on the surface of the virus-infected cells. We have previously demonstrated that the virus filaments formed at the distal end of cell surface projections that are composed of the G protein, suggesting that the anti-F staining would more accurately define the virus filaments on the surface of infected cells[9]. We therefore examined the distribution of the G, F and M proteins on virus-infected cells since these are the major virus structural proteins that interact with the virus envelope and define the morphology of the virus filaments [57].

Mock-infected (SFig. 1A and B) and RSV-infected cells (SFig. 1C(i) and (ii)) were stained using anti-F and anti-G and examined using immunofluorescence (IF) microscopy and phase contrast (PC) microscopy. Antibody staining was only observed in the virus-infected cells, which confirmed the specificity of these immunological reagents in this current study. Representative non-permeabilised anti-G and anti F co-stained RSV-infected cells were examined in greater detail using confocal microscopy at a focal plane that allowed imaging of the cell periphery (Fig. 3A) and cell top (Fig, 3B), and the presence of anti-F and anti-G co-stained virus filaments was detected at the cell periphery and cell top. The anti-F and anti-G co-stained virus filaments were also located at the distal ends of distinct anti-G-stained projections at both locations, but these different staining distributions were more noticeable at the cell periphery where the anti-G projections extended to the adjacent non-infected cells. These longer anti-G-stained projections that extended from the cell periphery were similar in appearance to filopodia, and in this current study we refer to these longer anti-G-stained projections as long filamentous projections (LFP); this is to distinguish them from the anti-G and anti-F co-stained virus filaments. These data indicated that while the anti-G antibody stained both the LFP and virus filaments, the anti-F staining was confined to the virus filaments, confirming our earlier observations [9]. While the virus filaments that form at these different locations are predicted to be infectious, it is presumed that more efficient localised virus cell-to-cell transmission may occur via the LFP since these structures come into direct contact with the surrounding non-infected cells. Collectively, these imagining data described above confirmed that anti-F staining could specifically delineate the virus particles that form on the surface of the RSV-infected A459 cells, and anti-F staining was primarily used to detect the virus filaments in the subsequent analyses.

**Figure 3.**
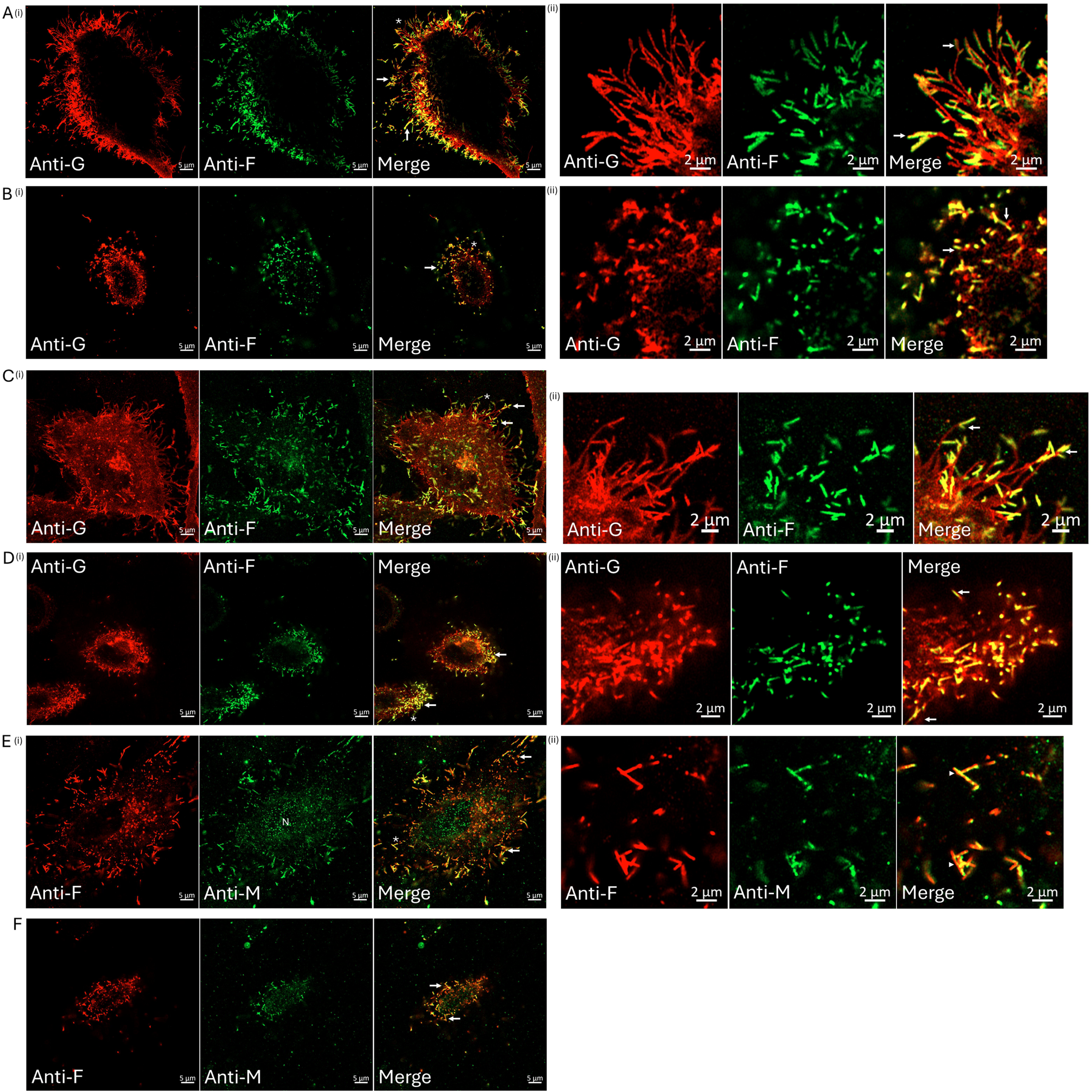
Distribution of the virus filaments on the surface of RSV-infected A549 cells. A549 cell monolayers were RSV-infected using a multiplicity of infection (moi) of 0.05 and at 24 hrs post-infection (hpi) the non-permeabilized cells were co-stained with anti-F and anti-G. Representative co-stained cell was examined using confocal microscopy and a series of images in the Z-plane (Z-stack) were recorded and images at **(A)** the cell periphery and **(B)** close to the cell top are shown. **(C to F)** The permeabilized cells were co-stained with **(C and D)** anti-F and anti-G and **(E) and (F)** anti-F and anti-M. Representative co-stained cell was examined using confocal microscopy and a series of images in the Z-plane (Z-stack) were recorded. Images at **(C)** and **(E)** the cell periphery and **(D)** and **(F)** close to the cell top are shown. In all plates **(A to E)** the plates marked as (ii) are enlarged images taken from the corresponding plate marked as (i) in the area highlighted (*) and the co-stained virus filaments are highlighted (white arrows). In the confocal microscopy images individual image channels and the merged images are shown.

We also examined infected cells using anti-M staining to confirm that the M protein was present in the virus filaments that were delineated by the anti-F-staining. In mock-infected and RSV-infected cells stained using anti-M and imaged using IF microscopy and PC microscopy (SFig. 1D) the antibody-staining was only observed in the RSV-infected cells, which confirmed the specificity of the anti-M antibody. We also examined anti-F and anti-M co-staining in RSV-infected cells that were non-permeabilized (SFig. 1E(i)) or permeabilised (SFig. 1E(ii)). This confirmed that the anti-M staining was only detected in the permeabilised cells, consistent with the M protein being located on the underlying surface of the virus envelope. Permeabilised RSV-infected cells were co-stained with anti-F and anti-G (Fig. 3C and D), and with anti-F and anti-M (Fig. 3E and F) and the individual staining patterns examined using confocal microscopy. The distribution of the F and G protein in representative permeabilised anti-G and anti F co-stained cells exhibited a similar distribution of the F and G protein to that described above in the non-permeabilised cells. Representative permeabilised anti-F and anti-M co-stained RSV-infected cells exhibited high levels of anti-F and anti-M co-staining in the virus filaments, indicating the presence of the M protein in the anti-F-stained virus filaments.

### 3. The distribution virus particles and the glycocalyx on the surface of RSV-infected cells

We examined the distribution of the glycocalyx in the context of the virus particles that formed on the surface of RSV-infected A549 cells. In this analysis we used WGA- AL488, anti-HS, and anti-SYND4 to visualise the cell glycocalyx on these cells. The virus filaments were detected on the surface of the infected cells using antibodies that recognise the virus structural proteins in the virus filaments. This involved mainly using anti-F staining, but we also used anti-G, anti-M and anti-N (that recognises the RSV NC) staining to detect the virus filaments as appropriate.

Representative non-permeabilised RSV-infected cells that were co-stained with WGA-AL488 and anti-G were examined at a focal plane that allowed imaging of the cell periphery (Fig. 4A) and cell top (Fig. 4B). In virus-infected cells WGA-AL488 and anti-G co-staining at the cell periphery was observed throughout the anti-G stained LFP, and on the virus filaments that form at the distal ends of the LFP and on the anti-G-stained virus filaments on the cell top. Non-permeabilised RSV-infected cells were also co-stained with WGA-AL488 and anti-F (Fig. 4C and D) and then examined using confocal microscopy to determine the respective distribution of the glycocalyx and virus particles. Representative non-permeabilised RSV-infected cells were also co-stained with WGA-AL488 and anti-F were examined at a focal plane that allowed imaging of the cell periphery (Fig. 4C) and cell top (Fig. 4D). In each case the WGA-AL488 and anti-F co-staining on the virus filaments that form at the cell periphery and on the cell top was noted.

**Figure 4.**
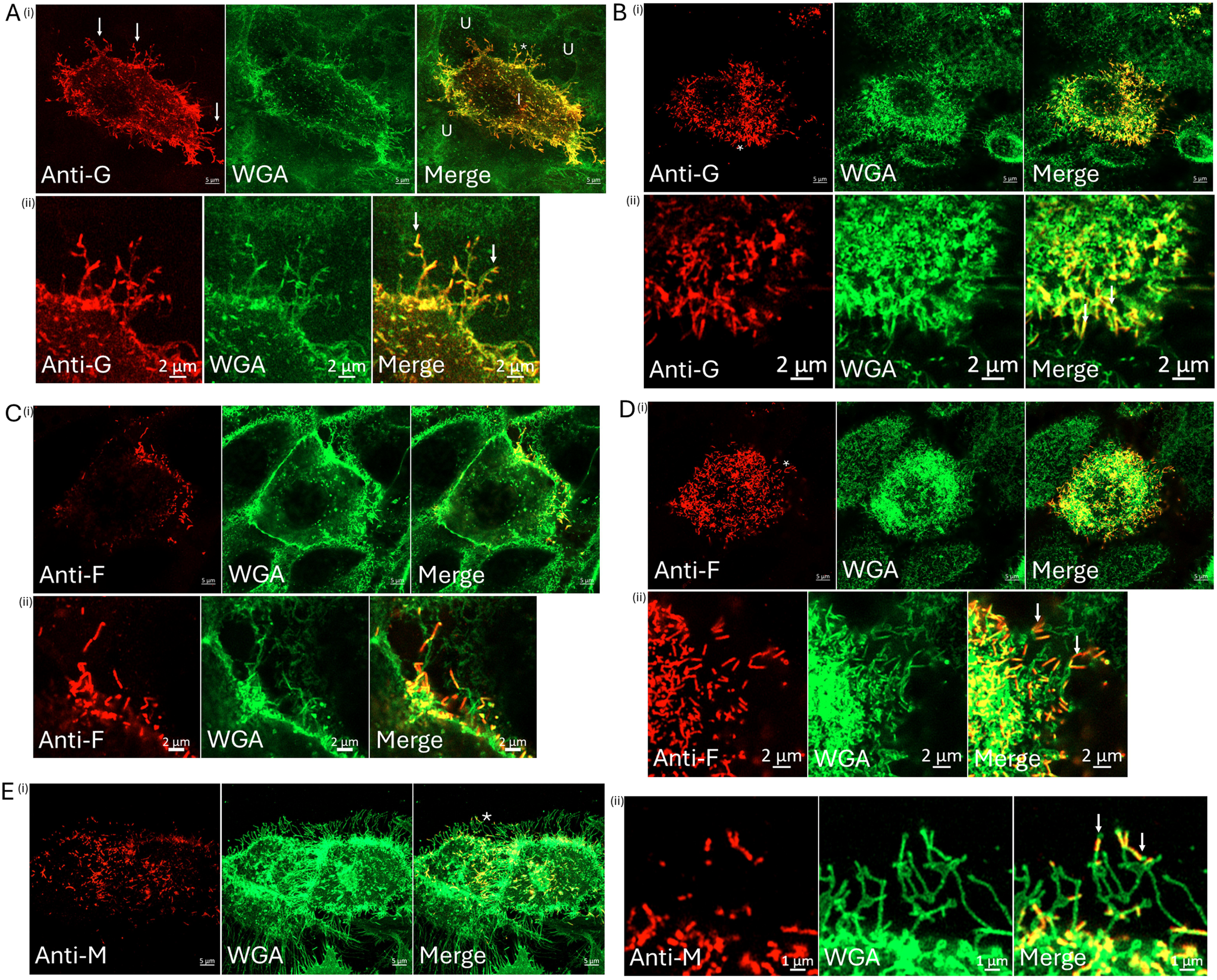
Distribution of the WGA-AL488-stained glycocalyx and virus filaments on the surface of RSV-infected A549 cells. A549 cell monolayers were RSV-infected using a multiplicity of infection (moi) of 0.05 and at 24 hrs post-infection (hpi). The non-permeabilised cells were co-stained using **(A and B)** WGA-AL488 and anti-G and **(C** and **D)** WGA-AL488 and anti-F. Representative co-stained cells were imaged using confocal microscopy and a series of images in the Z-plane (Z-stack) were recorded. Images at **(A and C)** the cell periphery and **(B and D)** close to the cell top from the image series are shown. In each case (ii) is an enlarged image from area highlighted (white *) in (i). **(E)** non-permeabilised RSV-infected cells were co-stained with WGA-AL488 and then the cells were permeabilised and stained with anti-M and representative co-stained cells were imaged using confocal microscopy. A series of images in the Z-plane (Z-stack) were recorded and an image from the cell periphery is shown **(E)** (ii) is an enlarged image from area highlighted (white *) in **(E)** (i). The co-stained virus filaments are highlighted (white arrows) and U highlights non-infected cells in the same field of view. In the confocal microscopy images individual image channels and the merged images are shown.

The M protein is also a major virus structural protein that co-stained the anti-F-stained virus filaments, and therefore we also examined the distribution of the M protein and WGA- AL488 staining on virus-infected cells. The M protein is distributed beneath the virus envelope, and the detection of the M protein in the virus filaments required cell permeabilization prior to antibody staining. Therefore, the RSV-infected cells were first surface stained using WGA-AL488, after which the cells were permeabilised and co-stained with anti-M (Fig. 4E(i)). Analysis of representative cells by confocal microscopy showed anti-M and WGA-AL488 co-staining in the virus filaments that form at the distal ends of the WGA-AL488-stained LFP and at the cell top (Fig. 4E(ii)). These data also indicated that like the F protein, the M protein was not present throughout the WGA-AL488-stained LFP, but it was only present in the virus filaments that form at the distal ends of the LFP.

Due to the low moi used in this analysis, the adjacent non-infected cells (indicated by e.g. the absence of anti-G and anti-F staining) were also detected in the same field of view. The WGA-AL488 staining on these cells appeared to be primarily located in filamentous projections whose appearance was consistent with microvilli as described in the above analysis on the mock-infected cells described above. The WGA-AL488 staining of the virus filaments provided evidence that the virus filaments were bound by the host cell glycocalyx. The virus glycoproteins are modified by N-linked and O-linked glycosylation, and it has been demonstrated that specific lectins that recognise these sugar structures associated with O-linked glycosylation also bind the G protein [58]. The WGA-AL488 probe recognises terminal N-acetyl-D-glucosamine and sialic acid moieties, and it is currently unclear to what extent the WGA-AL488 probe also binds the virus glycoproteins that are displayed on the surface of the virus filaments. However, when compared with the surrounding non-infected cells we generally noted a slight increase in the WGA-AL488-staining intensity on the virus filaments that form on the virus-infected cells. We assume that this apparently small increase in staining intensity may be due to the additional presence of the virus glycoproteins that are displayed on the surface of the virus envelope. This suggests that the virus glycoproteins that are displayed on the surface of the virus filaments may also contribute to the biological properties of overlying cellular glycocalyx that surrounds the virus particles.

Non-permeabilised mock-infected and RSV-infected cells were co-stained with anti-F and anti-HS and imaged using IF microscopy (Fig. 5A). While the anti-F staining was observed only on RSV-infected cells, high levels of anti-HS staining on the surface of both mock-infected and RSV-infected cells was observed. However, when the cells were imaged under identical camera settings (e.g. exposure times) no detectable difference in the anti-HS staining intensity on mock-infected and RSV-infected cells was detected, which suggested that virus infection did not lead to a significant increase in HS levels on the surface of these cells. Examination of these cells at higher magnification using IF microscopy (Fig. 5A) suggested some alignment of the anti-F and anti-HS staining patterns on the surface of virus-infected cells, and this was examined in more detail by using confocal microscopy. Representative anti-F and anti-HS co-stained cells were examined at a focal plane that allowed imaging of the cell periphery (Fig. 5B and C(i)) and cell top (Fig. 5D(i)). While anti-HS staining was observed throughout the cell surface of the RSV-infected cells, specific anti-HS staining on the surface of the virus filaments that form at both the cell periphery (Fig. 5C(ii)) and at the cell top (Fig 5D(ii)) was also observed. The series of images from the same representative the anti-HS and anti-F-co-stained non-permeabilised infected cell that was captured in the Z-projection was reconstructed into a 3-dimensional image (Fig. 5E). This also allowed the distribution of the virus filaments and the HS-stained glycocalyx to be imaged on the surface of these cells and further highlighted the anti-HS staining on the virus filaments.

**Figure 5.**
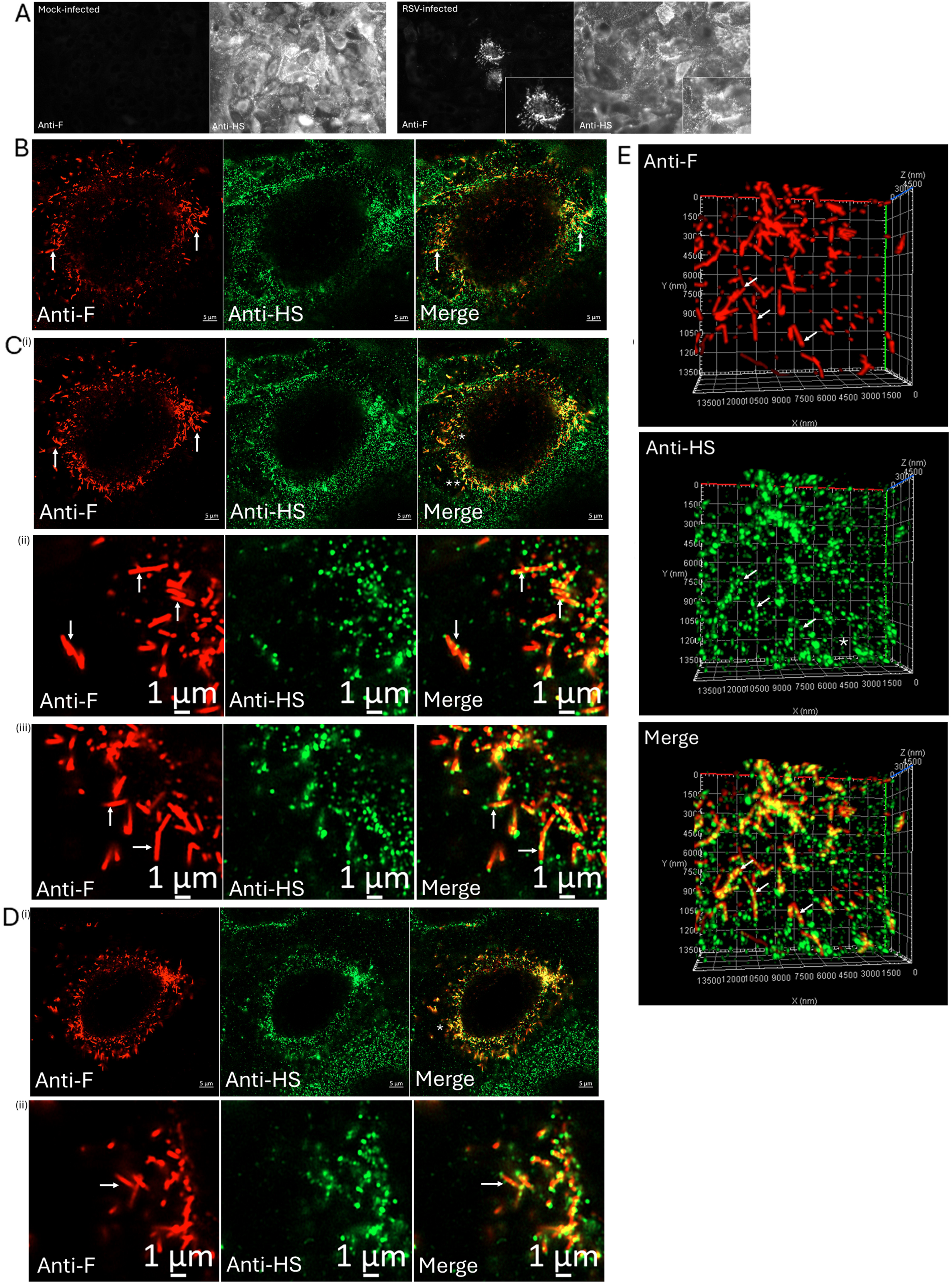
Distribution of the heparan sulphate and anti-F-stained virus filaments on the surface of RSV-infected A549 cells. **(A)** At 24 hrs post-infection mock-infected and RSV A549 cell monolayers were co-stained using anti-F and anti-heparan sulphate antibody (anti-HS) and the co-stained cells imaged using immunofluorescence microscopy (magnification x40 objective). Inset is an enlarged image taken the infected cell stained with each antibody. **(B-D)** A representative co-stained RSV-infected cell was examined using confocal microscopy. A series of images in the Z-plane (Z-stack) were recorded and images at **(B)** the cell periphery and **(C)** mid-section and **(D)** close to the cell top are shown. **(C)** (ii) and **(C)**(iii) are enlarged images taken from **(C)** (i) in the areas highlighted by * and ** respectively. **(D)** (ii) is an enlarged image taken from **(D)** (i) in the areas highlighted by *. The anti-F-stained virus filaments in the anti-F panel and the anti-F and anti-HS co-stained virus filaments in the merged images are highlighted (white arrows). **(E)** A series of images through an anti-F and anti-HS-co-stained infected cell was obtained in the Z projection and were processed into a single 3-dimentional image. The anti-F and anti-HS staining on the virus filaments are highlighted (white arrows). Also indicated is the anti-HS staining not associated with the virus filaments (*). In the confocal microscopy images individual image channels and the merged images are shown.

The anti-G and anti-F staining patterns on the virus filaments were continuous along the length of the virus filaments, and is consistent with our previous studies employing immuno-scanning electron microscopy that suggested close packing of the G protein on the virus filaments [59]. However, these antibody staining patterns contrasted with the localised and intermittent anti-HS staining pattern that was observed along the length of the virus filaments. Irrespective of these differences in the distribution of the anti-F and anti-HS staining patterns, these data demonstrate the presence of HS on the surface of the anti-F-stained virus filaments. This was consistent with the WGA-488 staining described above, providing additional evidence that the surface of virus particles was bounded by the glycocalyx. Since the anti-HS antibody recognition requires N-sulfation of GlcNAc residues on the HS chain, this suggested that the HS detected on the virus filaments was modified by GlcNAc N-sulfation. However, further sulfation at other sites in the repeating GlcA-GlcNAc disaccharide unit in the HS chains can occur by host cell sulfotransferases, and it is currently unclear to what extent the HS on the virus filaments was further modified by sulfation.

The RSV N protein is a major component of the virus NC, and the NC accumulate in the cytoplasm within the virus cytoplasmic inclusion bodies [13] and they are also packaged into the virus filaments that form on the surface of infected cells. We therefore also examined the distribution of HS and the virus filaments in RSV-infected cells that were co-stained with anti-N and anti-HS. Since the NC is packaged within the virus filament, the cells were first permeabilised prior to anti-N staining. Mock-infected and RSV-infected were permeabilised and co-stained using anti-N and imaging using IF microscopy and PC microscopy (SFig. 1F), and the anti-N staining was only observed in the RSV-infected cells which confirmed the specificity of the anti-N antibody. RSV-infected cells were permeabilised and co-stained with anti-N and anti-HS and representative co-stained cells were examined in detail using confocal microscopy. We have previously noted that in anti-N-stained RSV-infected cells the inclusion bodies stain more intensely when compared with the anti-N-stained virus filaments. This is presumably a reflection of the increased levels of the N protein that is present within the inclusion bodies compared with that in the virus filaments. Therefore, in this analysis the representative co-stained cells in the same field of view were imaged using two different camera exposure time settings, which allowed the anti-N-stained inclusion bodies and virus filaments to be more clearly defined in the same cell. At the lower camera exposure setting the individual anti-N-stained inclusion bodies were more clearly defined (Fig. 6A), and the absence of anti-HS staining in these structures was noted. Using a higher camera exposure setting the anti-N-stained virus filaments that form on these cells were better imaged (Fig. 6B(i)), which showed the presence of the anti-HS co-staining along the length of the anti-N-stained virus filaments (Fig. 6B(ii)). Although the anti-N-stained NC was packaged within the virus filaments, the anti-HS staining occurred along the length of the anti-N-stained virus filaments and was consistent with the surface staining obtained using the anti-F antibody described above.

**Figure 6.**
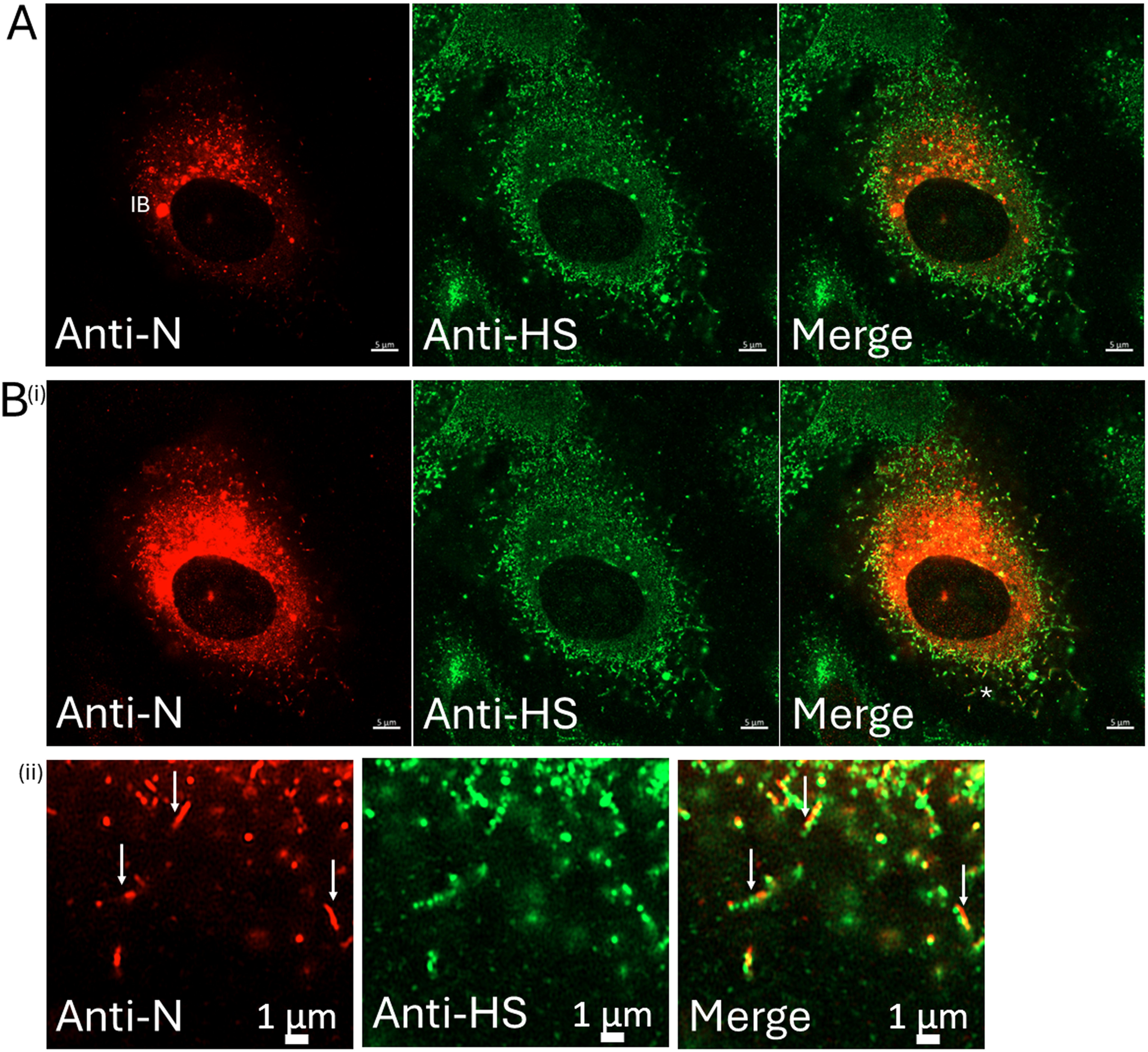
Distribution of the heparan sulphate and anti-N-stained virus filaments on the surface of RSV-infected A549 cells. A549 cell monolayers were RSV-infected and at 24 hrs post-infection (hpi) the permeabilized cells were co-stained with anti-N and anti-heparan sulphate antibody (anti-HS). A representative co-stained cell was examined using confocal microscopy. A series of images in the Z-plane (Z-stack) were recorded and an image at the cell periphery that shows the inclusion bodies and virus filaments is shown. **(A)** the anti-N staining was imaged at a shorter exposure and **(B)** a longer exposure. **(B)** (ii) is an enlarged image taken from **(B)** (i) in the areas highlighted by *. The inclusion bodies (IB) and co-stained virus filaments (white arrows) are highlighted. In the confocal microscopy images individual image channels and the merged images are shown.

We also examined the expression of the SYND4 protein on the surface of RSV-infected cells to visualise the glycocalyx in relation to the virus filaments. Non-permeabilised mock-infected and RSV-infected cells were co-stained with anti-F and anti-SYND4 and imaged using confocal microscopy. Representative co-stained mock-infected and RSV-infected cells were examined at a focal plane that allowed imaging of the virus filaments (Fig. 7). On the mock-infected cells a diffuse anti-SYND4 staining was noted (Fig 7A), and as expected anti-F staining was not detected. Although anti-SYND4 staining was observed throughout the surface of the RSV-infected cells, on these cells a proportion of the anti-SYND4 staining was detected on the surface of the anti-F-stained virus filaments (Fig. 7B and C(i)). Closer examination of the anti-F-labelled virus filaments confirmed the presence of the anti-SYND4 staining on the virus filaments (Fig. 7C(ii)), which exhibited a staining pattern that was similar in appearance to the anti-HS staining that was observed on the virus filaments. The series of images from the same representative anti-SYND4 and anti-F-co- stained calls obtained in the Z-projection was reconstructed into a 3-dimensional image (Fig. 7D). This also allowed the distribution of the virus filaments and the anti-SYND4 staining on the surface of these cells to be imaged and further highlighted the anti-HS staining on the virus filaments. Non-permeabilised RSV-infected cells were also co-stained with anti-G and anti-SYND4 and imaged using confocal microscopy, and representative co-stained cells were examined at a focal plane that allowed imaging of the virus filaments (Fig. 8). While anti- SYND4 staining was again observed throughout the surface of the RSV-infected cells, a proportion of the anti-SYND4 staining was detected on the surface of the anti-G-stained LFP at the periphery of the infected cells (Fig 8A(i) and (ii))) and was consistent with the imaging using the WGA-AL488 probe described above. In addition, anti-SYND4 staining was detected on the surface of the anti-G-stained virus filaments (Fig 8B(i),(ii) and (iii))), and these exhibited a similar staining pattern to that described above on the cells co-stained with anti-F and anti-SYND4 (Fig. 7B and C(ii)). These data indicated the presence of SYND4 protein on the surface of the virus filaments, and this provided additional evidence for the presence of the glycocalyx on the surface of the virus filaments that form on the surface of virus-infected cells.

**Figure 7.**
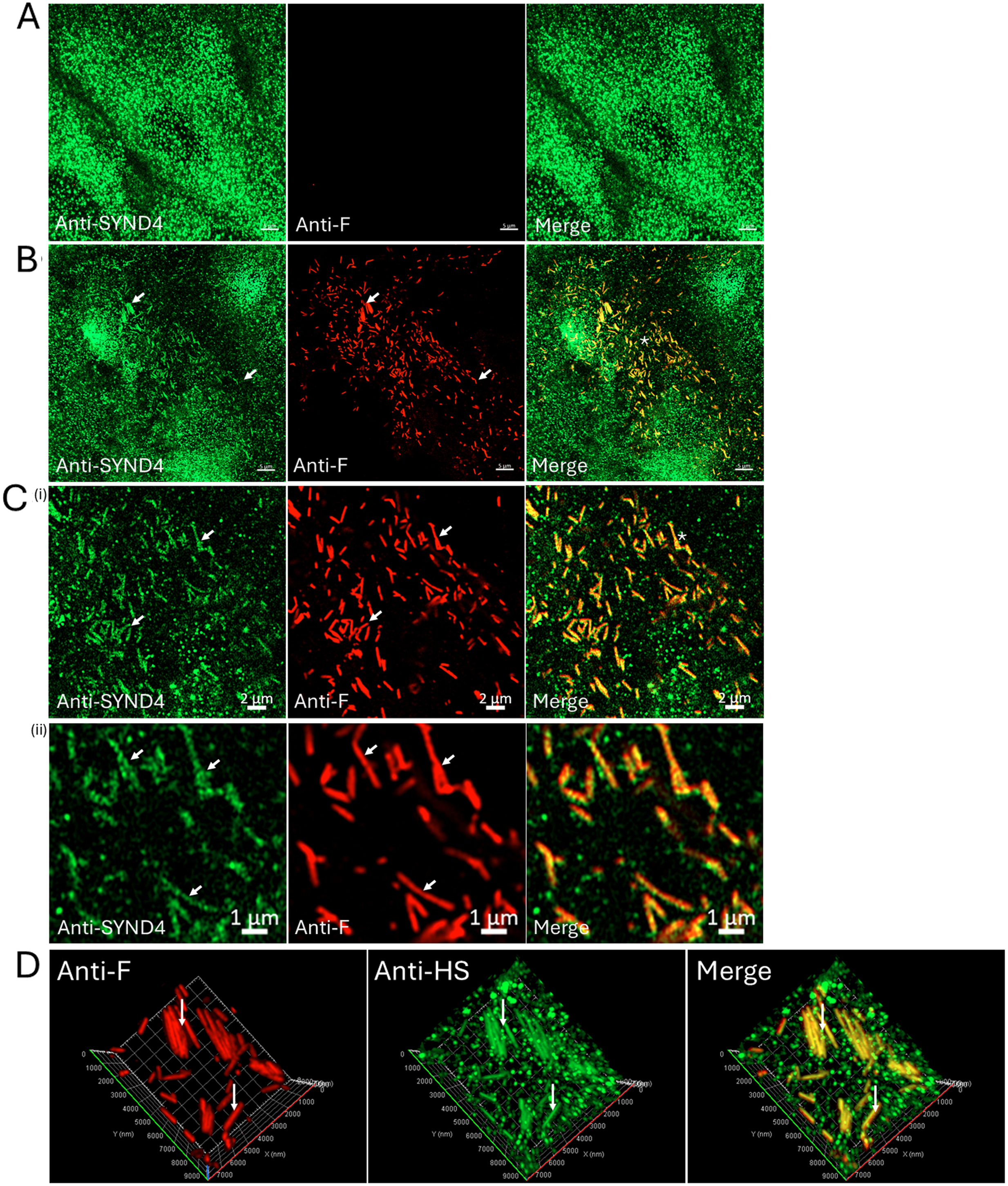
Distribution of the Syndecan-4 protein and anti-F-stained virus filaments on the surface of RSV-infected A549 cells. Mock-infected and RSV-infected A549 cell monolayers were co-stained using anti-F and anti-Syndecan-4 antibody (anti-SYND4) and the co-stained cells imaged using confocal microscopy. A series of images in the Z-plane (Z-stack) were recorded and individual image channels and the merged images are shown. Representative co-stained **(A)** Mock-infected and **(B and C**) RSV-infected cells at a corresponding focal plane that allows imaging of the virus filaments are presented. **(C**(i)) is an enlarged image taken from plate **(B)** in the area highlighted by (*), and (**C**(ii)) is a further enlarged image taken from plate **(C**(i)) in the area highlighted by (*). The anti-F and anti-SYDN4 stained virus filaments are highlighted (white arrows). **(D)** A series of images through an anti-F and anti-SYND4-co-stained infected cell was obtained in the Z projection and were processed into a single 3-dimentional image. The anti-F and anti-SYND4 staining on the virus filaments are highlighted (white arrows). In the confocal microscopy images, individual image channels and the merged images are shown.

**Figure 8.**
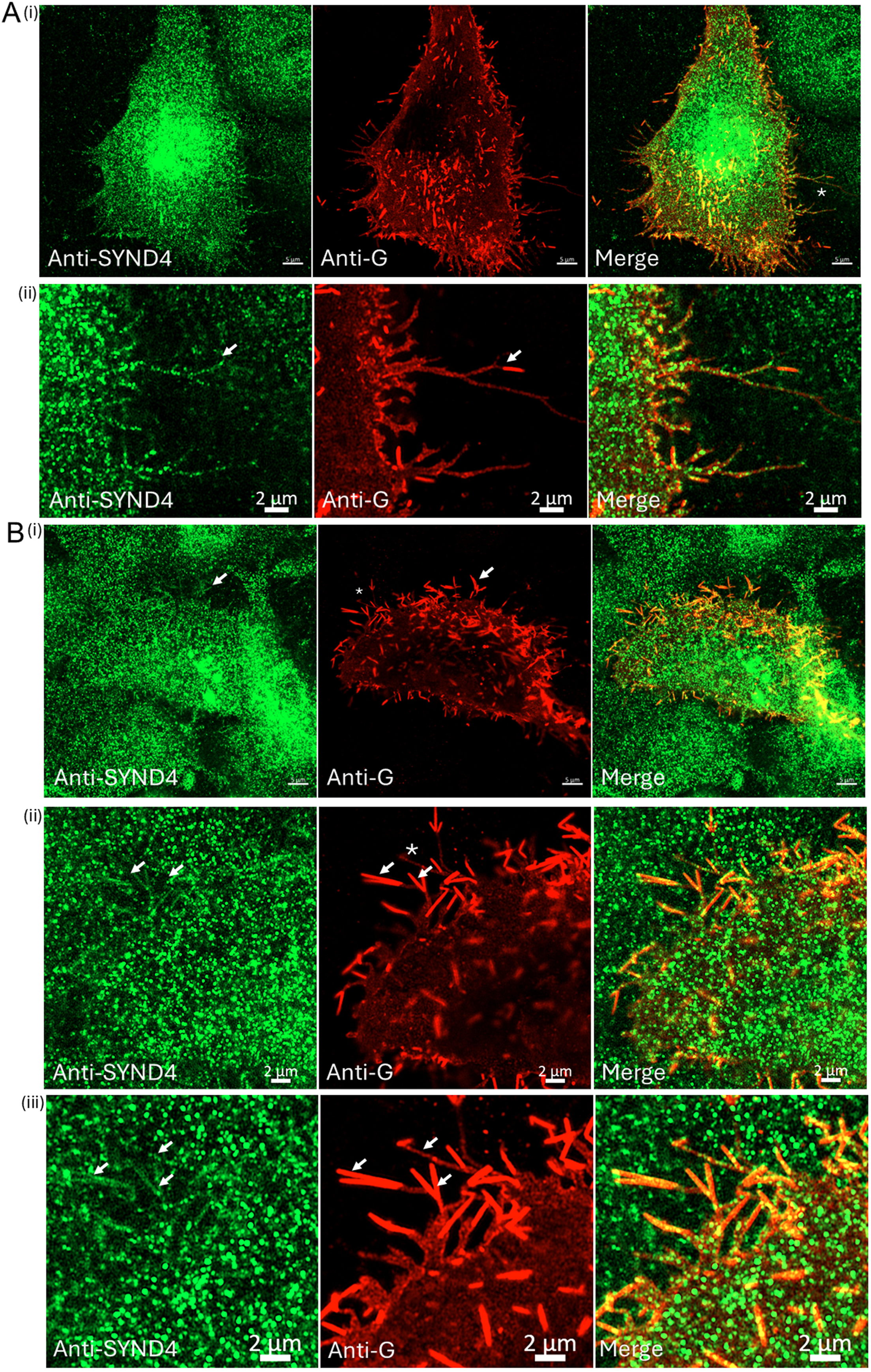
Distribution of the Syndecan-4 protein and anti-G-stained virus filaments on the surface of RSV-infected A549 cells. RSV-infected A549 cell monolayers were co-stained using anti-G and anti-Syndecan-4 antibody (anti-SYND4) and the co-stained cells imaged using confocal microscopy. **(A** (i) **and B**(i)) are two representative co-stained RSV-infected cells and a series of images in the Z-plane (Z-stack) were recorded and individual image channels and the merged images are shown. (**A**(ii)) is an enlarged image taken from plate **(A**(i)) in the area highlighted by (*). The anti-G and anti-SYDN4 co-stained LFP is highlighted (white arrow). (**B**(ii)) is an enlarged image taken from plate **(B**(i)) in the area highlighted by (*). (**B**(iii)) is a further enlarged image taken from plate **(B**(ii)) in the area highlighted by (*). The anti-G and anti-SYDN4 co-stained virus filaments are highlighted (white arrows). In the confocal microscopy images individual image channels and the merged images are shown.

### 4. The glycocalyx envelopes the virus filaments during the later stages of infection

The imaging data indicated that anti-HS staining on the surface of the virus filaments at a time of infection when virus particle assembly was complete, providing evidence that the virus filaments are bounded by the glycocalyx. In a further analysis we examined RSV-infected cells at a later stage of infection when significant levels of virus cell-to-cell transmission would be expected [9, 18]. Under the experimental conditions used in this study a significant level of virus cell-to-cell transmission at 48 hpi would be expected, and this was confirmed by examining cell monolayers at 24 and 48 hpi. The cell monolayers were co-stained using anti-F and either anti-G or anti-M and imaged using IF and PC microscopy (SFig. 2). While mainly single co-stained cells in the cell monolayer were detected at 24 hpi, by 48 hpi the presence of larger infected cell clusters predominated in the cell monolayer and indicated localised virus transmission. At 48 hpi the non-permeabilised cells were co-stained using anti-F and anti-HS and representative co-stained cells examined using confocal microscopy at a focal plane that allowed imaging of the cell periphery (Fig. 9A(i)) and cell top (Fig. 9B(i), C(i) and D). This analysis indicated increased levels of anti-HS staining on the anti-F-stained virus filaments, both at the cell periphery (Fig. 9A(ii)) and on the cell top (Fig. 9B(ii) and 8C(ii)). Imaging at the cell periphery also showed the presence of co-stained virus filaments on infected cells that made direct contact with the adjacent non-infected cells (that showed the absence of anti-F staining) in the cell monolayer (Fig. 9A(ii)). These data indicated that the HS (and the cell glycocalyx) was present on the virus filaments during the later stages of virus infection when significant levels of virus transmission had occurred, suggesting that the glycocalyx is a stable feature of the virus particles that form on virus-infected cells.

**Figure 9.**
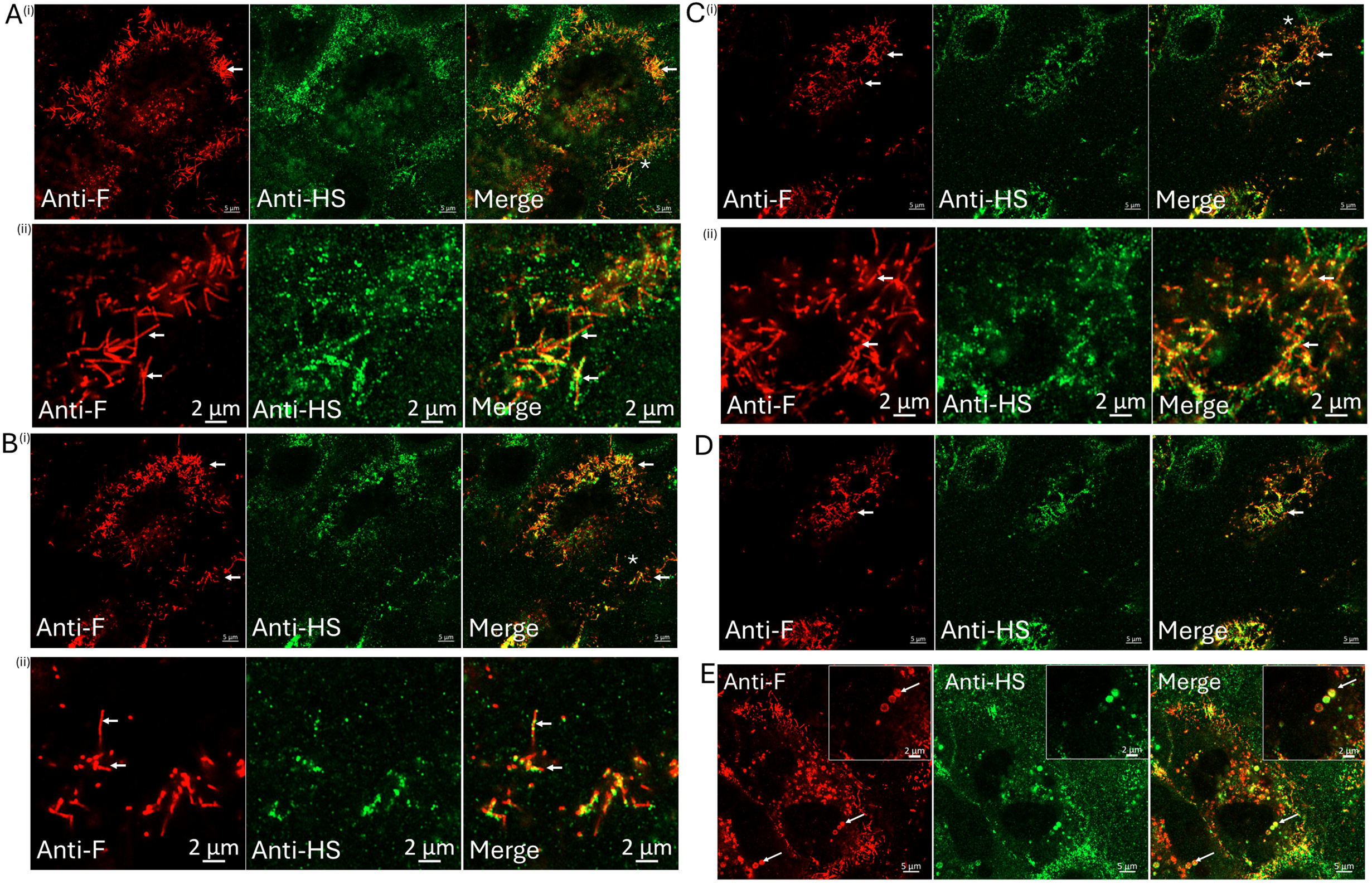
Distribution of the heparan sulphate and anti-F-stained virus filaments on the surface of RSV-infected A549 cells at 48 hrs post-infection. A549 cell monolayers were RSV-infected using a multiplicity of infection (moi) of 0.05 and at 48 hrs post-infection (hpi). A549 cell monolayers were co-stained using anti-F and anti-heparan sulphate antibody (anti-HS). A representative co-stained RSV-infected cells was examined using confocal microscopy. A series of images in the Z-plane (Z-stack) were recorded and images at **(A)** the cell periphery and **(B)** mid-section and **(C and D)** close to the cell top are shown. In plates **(A to C)** (ii) are enlarged images taken from (i) in the areas highlighted by *. The anti-F-stained virus filaments in the anti-F-stained image and the anti-F and anti-HS co-stained virus filaments in the merged image are highlighted (white arrows). **(E)** is an image showing the spherical virus particles highlighted by white arrows. Insets are enlarged images taken from the areas highlighted by *. In the confocal microscopy images individual image channels and the merged images are shown.

We have also previously reported that at the later stages of infection the appearance of low levels of virus particles that exhibit a spherical shaped morphology [9]. We had earlier hypothesised that the appearance of these virus structures may be related to the low levels of cell-free virus infectivity that is detected at the later stages of a multiple-cycle RSV infection resulting from cell detachment of the virus particles [18]. In this current analysis we also observed the presence of anti-F and anti-HS-stained spherical virus particles at this later time of infection (Fig. 9E), providing evidence that the HS was also associated with these virus structures. This further suggested that the glycocalyx may also be a feature of cell-free RSV particles that exhibit a pleiomorphic morphology after the detachment of the virus from the infected cells occurs.

### 5. The virus G protein is trafficked into cell surface projections that are bounded by the glycocalyx

The expression of the recombinant G protein in mammalian cells can form virus-like particles (VLPs), and that the co-expression of the recombinant F and G proteins can lead to the formation of VLP structures in which the F protein is stabilised in its pre-fusion form [60]. On cells expressing only the recombinant G protein, the appearance of anti-G-stained filamentous projections that are similar in appearance to virus filaments are detected [19]. These filamentous projections also contain several host cell factors that have been identified in virus particles that form on RSV-infected cells [19, 45], and we had earlier proposed that these filamentous projections may demarcate the sites of RSV particle assembly on these cells. In a final analysis we therefore used recombinant G protein expression in mammalian to examine if the glycocalyx was present at these sites of virus assembly prior to infection.

We first confirmed our earlier observation on the distribution of the recombinant G protein by comparing the expression patterns of the recombinant G protein with that of the recombinant F and M proteins. HEp2 cells were transfected with pCAGGS/G, pCAGGS/F and pCAGGS/M that express the recombinant G, F and M proteins respectively, and at 18 hrs post-transfection the cells were stained with anti-G, anti-F, and anti-M respectively, and imaged using IF microscopy (SFig 3A) and confocal microscopy (SFig. 3B-D). In non-permeabilised cells the recombinant G protein exhibited a prominent anti-G filamentous staining pattern, while the recombinant F protein exhibited a punctate anti-F staining pattern. The recombinant M protein exhibited a diffuse and uniform anti-M staining pattern in permeabilised cells. Immunoblotting of cell lysates prepared from these cells with the appropriate antibody showed protein species of the expected sizes for the respective virus protein (SFig. 3E). These data confirmed our earlier observations on recombinant G protein expression and highlighted the distinct cellular distribution of the recombinant expressed G protein compared with that exhibited by the recombinant expressed F and M proteins. Although we propose that these anti-G-stained protections delineate the sites of virus assembly, these are not the same as the virus filaments that form on virus-infected cells. During RSV infection the activation of specific cellular signalling pathways occurs that is required to modulate F-actin structures during virus particle assembly [15, 16, 19, 25], and in our hands these pathways were not activated in mock-infected cells expressing the recombinant G protein (unpublished observations).

Since the RSV infection work described was performed in A549 cells, in this analysis we examined the expression of the recombinant G protein in A549 cells. Although we used WGA-AL488 to detect the glycocalyx on the transfected cells, the co-staining of the WGA- AL488 and anti-HS in the mock-infected A549 cells suggested that WGA-AL488 staining in the transfected cells would also indicate the presence of other glycocalyx-associated factors e.g. HS. Non-permeabilised RSV-infected A549 cells were stained with WGA-AL488 and co-stained with anti-G and examined by confocal microscopy (Fig. 10 A and B), which confirmed the WGA-AL488 and anti-G co-staining throughout the anti-G stained LFP, and on the virus filaments that form at the distal ends of the LFP (Fig. 10A) and on the anti-G-stained virus filaments on the cell top (Fig. 10B). The A549 cells were transfected with pCAGGS/G, and after 18 hrs post-transfection the non-permeabilised cells were co-stained using anti-G and WGA-AL488. Imaging of anti-G stained pCAGGS/G-transfected cells using IF microscopy at higher magnification (Fig. 10C) showed that the recombinant expressed G protein also exhibited a prominent filamentous staining pattern in A549 cells. The imaging analysis also suggested an alignment between the filamentous anti-G staining pattern and the WGA-AL488-stained filamentous projections, and the co-stained cells were examined in more detail by using confocal microscopy. Representative anti-G and WGA- AL488 co-stained non-permeabilised transfected cells expressing the recombinant G protein were examined using confocal microscopy at a focal plane that allowed imaging of the cell periphery (Fig. 10D(i)) and (ii) mid-cell (Fig. 10E(i) and (ii)) and the cell top (Fig. 10F). In this analysis anti-G staining was observed in the WGA-AL488-stained filopodia (Fig. 10D(ii)) that were similar in appearance to the LFP that were detected in RSV-infected cells that were described above, and anti-G staining was also observed on other WGA-AL488- stained cell surface projections such as microvilli (Fig. 10E(ii) and F). The WGA-AL488- stained projections detected on cells expressing the recombinant G protein also exhibited a similar morphological appearance to the WGA-AL488-stained projections on the adjacent non-transfected cells (that were not stained with anti-G antibody) but that were visible in the same field of view. This suggested that the recombinant G protein was trafficked into existing cell surface projections that were bounded by the glycocalyx, rather than the expression of the G protein inducing the formation of novel filamentous structures. These data further suggested that the trafficking of the G protein into these preexisting cellular projections bounded by the glycocalyx during virus infection may facilitate virus transmission in virus-infected cells e.g. via the LFP in RSV-infected cells. In this context, stable interactions between the G protein and the other virus proteins that associated with the virus envelope (the F, SH and M proteins) have been described in RSV-infected cells [8, 61], suggesting that their interaction with the G protein may facilitate the recruitment these virus proteins into the LFP during the process of virus particle assembly.

**Figure 10.**
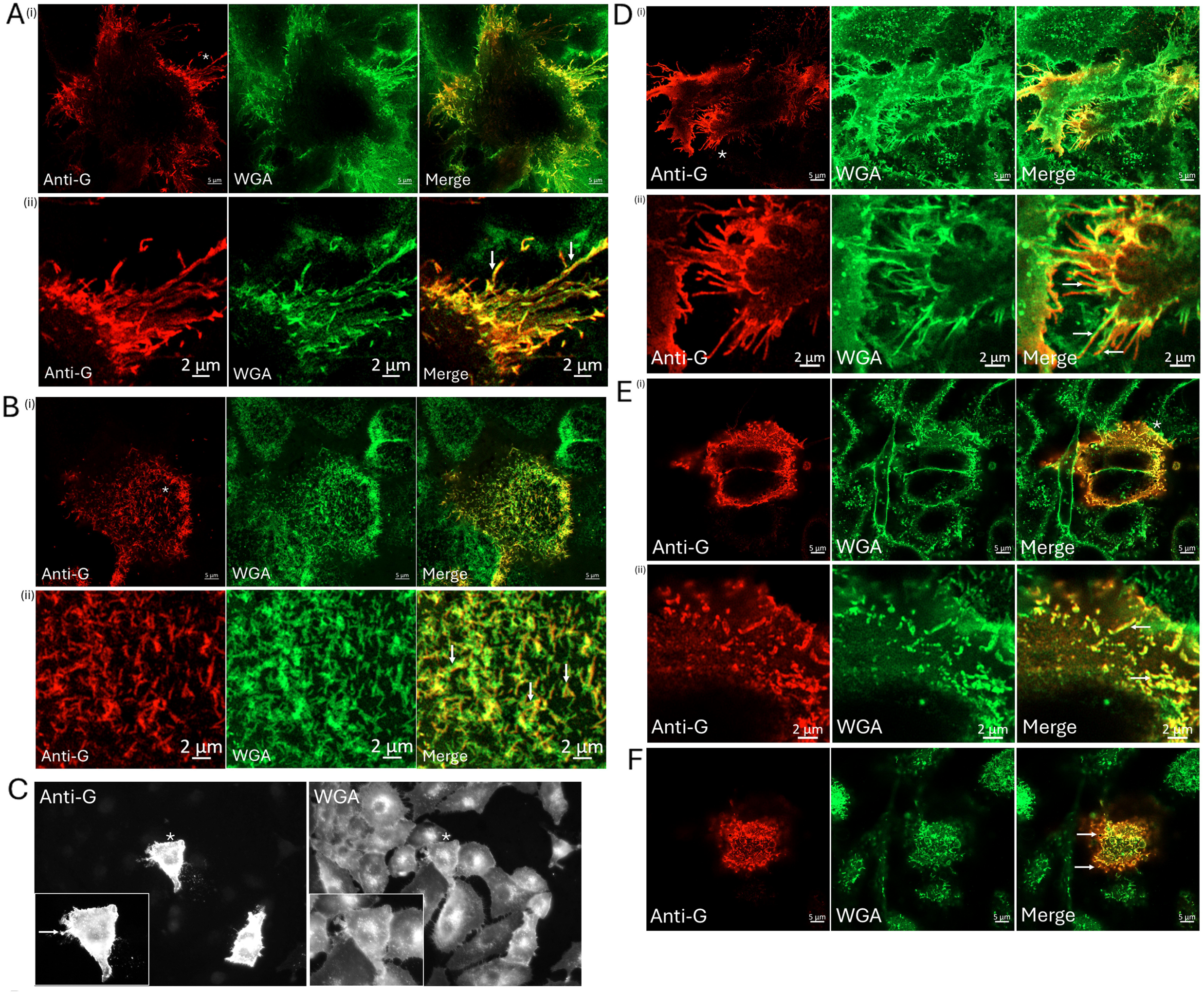
Distribution of the G protein the WGA-AL488-stained glycocalyx on the surface of A549 cells infected with RSV and expressing the recombinant G protein. A549 cell monolayers were RSV-infected using a multiplicity of infection (moi) of 0.05 and at 24 hrs post-infection (hpi). The non-permeabilised cells were co-stained with **(A and B)** WGA-AL488 and anti-G and representative co-stained cells were imaged using confocal microscopy and a series of images in the Z-plane (Z-stack) were recorded. Images at **(A)** the cell periphery and **(B)** close to the cell top from the image series are shown. In each case (ii) is an enlarged image from area highlighted (white *) in (i). **(C-F)** A549 cell monolayers were transfected with pCAGGS/G and after 18 hrs post-transfection **(C)** the cells were co-stained using anti-G and WGA-AL488 and the anti-G and WGA-AL488 co-stained pCAGGS/G-transfected cells were imaged using IF microscopy (magnification x40 objective). Inset is a magnified image of a representative anti-G and WGA-AL488 co-stained pCAGGS/G-transfected cell, highlighted by (*) in the main plate. **(D-F)** Representative anti-G and WGA-AL488 co-stained pCAGGS/G-transfected cells were examined using confocal microscopy. A series of images in the Z-plane (Z-stack) were recorded and images at **(D)** the cell periphery and **(E)** mid-section and **(F)** close to the cell top are shown. **(D)** (ii) is an enlarged images taken from **(D)** (i) and **(E)** (ii) is an enlarged image taken from **(E)** (i) in the areas highlighted by * in each respective plate. In the confocal microscopy images individual image channels and the merged images are shown.

## 6. Conclusion

Although previous studies have employed transmission electron microscopy (TEM) to examine RSV particle assembly on cells, these studies have not definitely demonstrated the presence of a well-developed glycocalyx on the surface of virus filaments. The reason for this is unclear and it is possible that the physical dimensions of the glycocalyx covering the virus filaments on virus-infected cells may be too small to be clearly defined and distinguished from the virus envelope. Specific experimental staining protocols are required to stabilise and visualise the glycocalyx for imaging by TEM in a vacuum [62, 63], and the glycocalyx may not be stable under the sample preparation protocols used in these previous studies. In this current study we have used an alternative experimental approach that used glycocalyx markers and light microscopy to image the virus-infected cells under close-to physiological conditions. Although our approach provides lower resolution image data when compared with electron microscopy, this approach allowed the detection of the glycocalyx on these cells and provided evidence for the presence of a well-developed glycocalyx on the surface of virus particles.

Our data provides the first evidence that during virus egress from the infected cell the virus filaments are bounded by the cell glycocalyx, suggesting that the virus filaments circumvent the glycocalyx by becoming enveloped by it. Since the glycocalyx can generate sufficient forces on the cell surface that can induce the formation of filamentous cellular structures such as filopodia and microvilli [39], its presence at the site of RSV particle assembly and on the RSV particles suggests that the glycocalyx may play an indirect role in defining the filamentous morphology of the virus filaments. We can also hypothesise that the glycocalyx may be stabilised on the surface of the virus filaments, and that this may involve the interaction between the surface-expressed RSV glycoproteins and the glycocalyx, by e.g. glycan-glycan interactions. In this context, a role for the glycocalyx in promoting HIV-1 cell entry via glycan-glycan interactions involving the HIV-1 glycoproteins has recently been presented [64].

We can assume that the surrounding glycocalyx may provide a protective barrier on the surface of the virus envelope that protects the integrity of the virus envelope from a range of harmful molecules that are expected to be produced by the cells during virus infection. However, the virus-associated glycocalyx may also play a more specific role in the biology of the RSV than just providing a protective barrier, and this will require further investigation. Interestingly the HS that is derived from the overlying glycocalyx was detected on the surface of the newly formed virus filaments on virus-infected cells. The inhibitory effects of adding HS to RSV during the initial stages of virus cell adsorption is established and has provided evidence for an established role for HS binding in facilitating RSV cell attachment [43, 44, 65]. However, HS has also been proposed to play a role in the cell attachment for an array of different virus with different tissue tropisms, and some viruses isolates can even be tissue culture adapted to use HS as a receptor during cell entry[66, 67]. In addition, in differentiated human airway epithelial cultures systems the RSV attaches to ciliated cells via the cell receptor CX3CR1 and in a HS-independent manner [4]. Collectively these observations suggest that HS may play a role in the virus replication cycle that is distinct from that envisaged during a typical interaction between a virus attachment protein and its virus host cell receptor. In this context in the binding of virus to HS on the cell surface has been proposed as a general strategy used by viruses that enable the closer interaction of the virus with the cell surface prior to virus entry [68]. This also suggests that the HS on the envelope of virus filaments may facilitate the interaction of RSV with the host cell plasma membrane during the early stages of infection. The plasma membrane is formed by a phospholipid bilayer, and an electrical potential is maintained across its boundary, with the inner leaflet in the bilayer exhibiting a negative charge and the outer leaflet exhibiting a positive charge (Alberts B, Johnson A, Lewis J, et al. Molecular Biology of the Cell. 4th edition. New York: Garland Science; 2002). In this context, the HS moieties that are present on the virus filaments are also expected to possess a strong negative charge [69], and this may impart a negative charge on the surface of the virus filaments that could then interact with the positively charged outer leaflet of the plasma membrane. Such an event would be expected to allow a closer interaction of the virus with the cell surface during the initial stages of infection and prior to engagement with a cell receptor. In addition, the HS moieties that are displayed on the surface of the virus filaments could potentially also interact with HS-binding host cell factors that are expressed on the surface of the host cell, and this could also facilitate the attachment of the virus to the cell surface. The nucleolin and annexin-2 proteins have both been proposed to act as cell receptors for RSV in permissive cell types [70, 71], and in this context both nucleolin and annexin-2 are also established as HS-binding proteins [72, 73]. Earlier studies by Bourgeois and collegues had demonstrated that the pretreatment of infectious RSV with an antibody against HS was able to neutralise RSV infection in permissive cell lines [74]. This early study suggested that HS-like structures may be present on the surface of RSV particles, and it was proposed that these could facilitate engagement of the virus with the cell membrane. Our current observations are consistent with the previous findings described by Bourgeois and collegues and provide the first direct evidence for the display of HS on the surface of virus particles. In addition, the inhibitory effect of HS- addition during virus cell adsorption that has been reported by others and our findings are also consistent with these previous observations. The addition of exogenous HS to cells at the time of virus adsorption would be expected to reduce the efficiency of binding of the virus-associated HS to the host cell by competing for HS-binding sites on the cell surface.

It is currently unclear how the HS that is displayed on the surface of the virus particles is anchored in the glycocalyx and the identity of the protein to which the HS chains are attached is currently unknown. Although previous studies have suggested that the G and F protein can both bind HS [75], we observed differences in the respective anti-G and anti-F staining patterns when compared with the corresponding anti-HS staining patterns on the virus filaments. Furthermore, the anti-HS staining pattern on the virus filaments was similar in appearance to the corresponding anti-HS staining pattern that was present on the WGA- AL488-stained filamentous projections on mock-infected cells. This suggests that if the HS on the virus filaments was bound to a protein that it may involve a host cell protein that is present in the glycocalyx. In this context, the presence of HS-modified SYND4 protein on the surface of the virus filaments may also potentially be one of the sources of the HS that we detected on the virus filaments. However, although the SYND4 protein is normally embedded in the plasma membrane via its transmembrane domain, under certain physiological conditions the HS modified ectodomain of the SYND4 protein can be excised by cellular activity [76, 77]. It is therefore not clear to what extent the population of the SYND4 protein that is detected in glycocalyx on the virus filaments is inserted into the virus envelope or if it is just contained within the overlying glycocalyx matrix. However, nevertheless the SYND4 protein and its associated HS is displayed on the surface of the virus filaments. Interestingly, the presence of the SYND4 protein on the virus filament may also have other consequences in the biology of the RSV. The SYND4 protein is also known to interact with the F-actin network and its associated signalling molecules [78, 79] and that these host cell factors play a role in RSV particle assembly and transmission[15, 19, 26].

Although we have examined the distribution of HS and the SYND4 protein on the surface of these cells, the glycocalyx on A549 cells is expected to contain other specific cellular components. These may be relevant to the biology of the RSV, and this will require further investigation. While future research will be required to better define the composition of the glycocalyx that envelopes the RSV particles, this will be challenging due to its expected biochemical complexity and the technical difficulties of examining the glycocalyx [62]. Interestingly, in respiratory tissue the SYND4 protein is highly expressed in airway epithelial cells and alveolar cells (https://www.proteinatlas.org/ENSG00000124145-SDC4/tissue), and its involvement in lung injury due to inflammation has been reported [35, 80]. Therefore, future work will also be required to determine if the presence of the glycocalyx (and its associated cellular factors) on the virus envelope has a clinical context during RSV infection, which may open new avenues for antiviral research. Lastly, our data suggests that the glycocalyx that is derived from the infected host cell is an integral part of the virus particle structure and suggests that this cellular structure should be included in structural models that are used to describe the mature RSV particles.

## Acknowledgements

This research received no specific grant from any funding agency in the public, private or not-for-profit commercial sectors.

## References

1. Nair, H., et al., Global burden of acute lower respiratory infections due to respiratory syncytial virus in young children: a systematic review and meta-analysis. Lancet, 2010. 375(9725): p. 1545-55.

2. Roberts, S.R., R.W. Compans, and G.W. Wertz, Respiratory syncytial virus matures at the apical surfaces of polarized epithelial cells. J Virol, 1995. 69(4): p. 2667–73.

3. Levine, S., R. Klaiber-Franco, and P.R. Paradiso, Demonstration that glycoprotein G is the attachment protein of respiratory syncytial virus. J Gen Virol, 1987. 68 **( Pt** **9**): p. 2521–4.

4. Johnson, S.M., et al., Respiratory Syncytial Virus Uses CX3CR1 as a Receptor on Primary Human Airway Epithelial Cultures. PLoS Pathog, 2015. 11(12): p. e1005318.

5. Sugrue, R.J., et al., Furin cleavage of the respiratory syncytial virus fusion protein is not a requirement for its transport to the surface of virus-infected cells. J Gen Virol, 2001. 82(Pt 6): p. 1375–86.

6. Gonzalez-Reyes, L., et al., Cleavage of the human respiratory syncytial virus fusion protein at two distinct sites is required for activation of membrane fusion. Proc Natl Acad Sci U S A, 2001. 98(17): p. 9859–64.

7. Zimmer, G., K.K. Conzelmann, and G. Herrler, Cleavage at the furin consensus sequence RAR/KR(109) and presence of the intervening peptide of the respiratory syncytial virus fusion protein are dispensable for virus replication in cell culture. J Virol, 2002. 76(18): p. 9218–24.

8. Low, K.W., et al., The RSV F and G glycoproteins interact to form a complex on the surface of infected cells. Biochem Biophys Res Commun, 2008. 366(2): p. 308–13.

9. Huong, T.N., et al., Evidence that an interaction between the respiratory syncytial virus F and G proteins at the distal ends of virus filaments mediates efficient multiple cycle infection. Virology, 2024. 591: p. 109985.

10. Grosfeld, H., M.G. Hill, and P.L. Collins, RNA replication by respiratory syncytial virus (RSV) is directed by the N, P, and L proteins; transcription also occurs under these conditions but requires RSV superinfection for efficient synthesis of full-length mRNA. J Virol, 1995. 69(9): p. 5677–86.

11. Yu, Q., R.W. Hardy, and G.W. Wertz, Functional cDNA clones of the human respiratory syncytial (RS) virus N, P, and L proteins support replication of RS virus genomic RNA analogs and define minimal trans-acting requirements for RNA replication. J Virol, 1995. 69(4): p. 2412–9.

12. Collins, P.L., et al., Transcription elongation factor of respiratory syncytial virus, a nonsegmented negative-strand RNA virus. Proc Natl Acad Sci U S A, 1996. 93(1): p. 81–5.

13. Garcia, J., et al., Cytoplasmic inclusions of respiratory syncytial virus-infected cells: formation of inclusion bodies in transfected cells that coexpress the nucleoprotein, the phosphoprotein, and the 22K protein. Virology, 1993. 195(1): p. 243–7.

14. Huong, T.N., et al., A sustained antiviral host response in respiratory syncytial virus infected human nasal epithelium does not prevent progeny virus production. Virology, 2018. 521: p. 20–32.

15. Gower, T.L., et al., RhoA signaling is required for respiratory syncytial virus-induced syncytium formation and filamentous virion morphology. J Virol, 2005. 79(9): p. 5326–36.

16. Ravi, L.I., et al., Increased hydroxymethylglutaryl coenzyme A reductase activity during respiratory syncytial virus infection mediates actin dependent inter-cellular virus transmission. Antiviral Res, 2013. 100(1): p. 259–68.

17. Mehedi, M., et al., Actin-Related Protein 2 (ARP2) and Virus-Induced Filopodia Facilitate Human Respiratory Syncytial Virus Spread. PLoS Pathog, 2016. 12(12): p. e1006062.

18. Huong, T.N., et al., Evidence for a biphasic mode of respiratory syncytial virus transmission in permissive HEp2 cell monolayers. Virol J, 2016. 13: p. 12.

19. Ravi, L.I., et al., Virus-induced activation of the rac1 protein at the site of respiratory syncytial virus assembly is a requirement for virus particle assembly on infected cells. Virology, 2021. 557: p. 86–99.

20. Brown, G., et al., Analysis of the interaction between respiratory syncytial virus and lipid-rafts in Hep2 cells during infection. Virology, 2004. 327(2): p. 175–85.

21. Brown, G., H.W.M. Rixon, and R.J. Sugrue, Respiratory syncytial virus assembly occurs in GM1-rich regions of the host-cell membrane and alters the cellular distribution of tyrosine phosphorylated caveolin-1. J Gen Virol, 2002. 83(Pt 8): p. 1841–1850.

22. Ludwig, A., et al., Caveolae provide a specialized membrane environment for respiratory syncytial virus assembly. J Cell Sci, 2017. 130(6): p. 1037–1050.

23. McCurdy, L.H. and B.S. Graham, Role of plasma membrane lipid microdomains in respiratory syncytial virus filament formation. J Virol, 2003. 77(3): p. 1747–56.

24. Brown, G., et al., Caveolin-1 is incorporated into mature respiratory syncytial virus particles during virus assembly on the surface of virus-infected cells. J Gen Virol, 2002. 83(Pt 3): p. 611–621.

25. Malhi, M., et al., Statin-mediated disruption of Rho GTPase prenylation and activity inhibits respiratory syncytial virus infection. Commun Biol, 2021. 4(1): p. 1239.

26. Jeffree, C.E., et al., Ultrastructural analysis of the interaction between F-actin and respiratory syncytial virus during virus assembly. Virology, 2007. 369(2): p. 309–23.

27. Kanyo, N., et al., Single-cell adhesivity distribution of glycocalyx digested cancer cells from high spatial resolution label-free biosensor measurements. Matrix Biol Plus, 2022. 14: p. 100103.

28. Peter, B., et al., Glycocalyx Components Detune the Cellular Uptake of Gold Nanoparticles in a Size- and Charge-Dependent Manner. ACS Appl Bio Mater, 2023. 6(1): p. 64–73.

29. Richter, J.R. and R.D. Sanderson, The glycocalyx: Pathobiology and repair. Matrix Biol Plus, 2023. 17: p. 100128.

30. Mockl, L., et al., Quantitative Super-Resolution Microscopy of the Mammalian Glycocalyx. Dev Cell, 2019. 50(1): p. 57–72 e6.

31. Mockl, L., The Emerging Role of the Mammalian Glycocalyx in Functional Membrane Organization and Immune System Regulation. Front Cell Dev Biol, 2020. 8: p. 253.

32. Sun, W.W., et al., Nanoarchitecture and dynamics of the mouse enteric glycocalyx examined by freeze-etching electron tomography and intravital microscopy. Commun Biol, 2020. 3(1): p. 5.

33. Li, Q., et al., Characterization of Cell Glycocalyx with Mass Spectrometry Methods. Cells, 2019. 8(8).

34. Reitsma, S., et al., The endothelial glycocalyx: composition, functions, and visualization. Pflugers Arch, 2007. 454(3): p. 345–59.

35. Gopal, S., et al., Syndecan receptors: pericellular regulators in development and inflammatory disease. Open Biol, 2021. 11(2): p. 200377.

36. Gondelaud, F. and S. Ricard-Blum, Structures and interactions of syndecans. FEBS J, 2019. 286(15): p. 2994–3007.

37. Piplani, N., et al., Bulky glycocalyx shields cancer cells from invasion-associated stresses. Transl Oncol, 2024. 39: p. 101822.

38. Kuo, J.C. and M.J. Paszek, Glycocalyx Curving the Membrane: Forces Emerging from the Cell Exterior. Annu Rev Cell Dev Biol, 2021. 37: p. 257–283.

39. Shurer, C.R., et al., Physical Principles of Membrane Shape Regulation by the Glycocalyx. Cell, 2019. 177(7): p. 1757–1770 e21.

40. Huang, X., et al., Association between plasma glycocalyx component levels and poor prognosis in severe influenza type A (H1N1). Sci Rep, 2022. 12(1): p. 163.

41. Puerta-Guardo, H., D.R. Glasner, and E. Harris, Dengue Virus NS1 Disrupts the Endothelial Glycocalyx, Leading to Hyperpermeability. PLoS Pathog, 2016. 12(7): p. e1005738.

42. Tay, E.A., et al., Protecting the endothelial glycocalyx in COVID-19. PLoS Pathog, 2024. 20(5): p. e1012203.

43. Hallak, L.K., et al., Iduronic acid-containing glycosaminoglycans on target cells are required for efficient respiratory syncytial virus infection. Virology, 2000. 271(2): p. 264–75.

44. Hallak, L.K., S.A. Kwilas, and M.E. Peeples, Interaction between respiratory syncytial virus and glycosaminoglycans, including heparan sulfate. Methods Mol Biol, 2007. 379: p. 15–34.

45. Radhakrishnan, A., et al., Protein analysis of purified respiratory syncytial virus particles reveals an important role for heat shock protein 90 in virus particle assembly. Mol Cell Proteomics, 2010. 9(9): p. 1829–48.

46. McDonald, T.P., et al., Evidence that the respiratory syncytial virus polymerase complex associates with lipid rafts in virus-infected cells: a proteomic analysis. Virology, 2004. 330(1): p. 147–57.

47. Rixon, H.W., et al., The small hydrophobic (SH) protein accumulates within lipid-raft structures of the Golgi complex during respiratory syncytial virus infection. J Gen Virol, 2004. 85(Pt 5): p. 1153–65.

48. Zhang, L., et al., Wheat germ agglutinin-conjugated fluorescent pH sensors for visualizing proton fluxes. J Gen Physiol, 2020. 152(6).

49. Targosz-Korecka, M., et al., Endothelial glycocalyx shields the interaction of SARS-CoV-2 spike protein with ACE2 receptors. Sci Rep, 2021. 11(1): p. 12157.

50. Targosz-Korecka, M., et al., Metformin attenuates adhesion between cancer and endothelial cells in chronic hyperglycemia by recovery of the endothelial glycocalyx barrier. Biochim Biophys Acta Gen Subj, 2020. 1864(4): p. 129533.

51. Zullo, J.A., et al., Exocytosis of Endothelial Lysosome-Related Organelles Hair- Triggers a Patchy Loss of Glycocalyx at the Onset of Sepsis. Am J Pathol, 2016. 186(2): p. 248–58.

52. van den Born, J., et al., Novel heparan sulfate structures revealed by monoclonal antibodies. J Biol Chem, 2005. 280(21): p. 20516–23.

53. David, G., et al., Developmental changes in heparan sulfate expression: in situ detection with mAbs. J Cell Biol, 1992. 119(4): p. 961–75.

54. Talsma, D.T., et al., MASP-2 Is a Heparin-Binding Protease; Identification of Blocking Oligosaccharides. Front Immunol, 2020. 11: p. 732.

55. van den Born, J., et al., Presence of N-unsubstituted glucosamine units in native heparan sulfate revealed by a monoclonal antibody. J Biol Chem, 1995. 270(52): p. 31303–9.

56. Cao, C., et al., Cholesterol-induced LRP3 downregulation promotes cartilage degeneration in osteoarthritis by targeting Syndecan-4. Nat Commun, 2022. 13(1): p. 7139.

57. Liljeroos, L., et al., Architecture of respiratory syncytial virus revealed by electron cryotomography. Proc Natl Acad Sci U S A, 2013. 110(27): p. 11133–8.

58. Collins, P.L. and G. Mottet, Oligomerization and post-translational processing of glycoprotein G of human respiratory syncytial virus: altered O-glycosylation in the presence of brefeldin A. J Gen Virol, 1992. 73 **( Pt** **4**): p. 849–63.

59. Jeffree, C.E., et al., Distribution of the attachment (G) glycoprotein and GM1 within the envelope of mature respiratory syncytial virus filaments revealed using field emission scanning electron microscopy. Virology, 2003. 306(2): p. 254–67.

60. McGinnes Cullen, L., et al., The Respiratory Syncytial Virus (RSV) G Protein Enhances the Immune Responses to the RSV F Protein in an Enveloped Virus-Like Particle Vaccine Candidate. J Virol, 2023. 97(1): p. e0190022.

61. Ghildyal, R., et al., Interaction between the respiratory syncytial virus G glycoprotein cytoplasmic domain and the matrix protein. J Gen Virol, 2005. 86(Pt 7): p. 1879–1884.

62. Haymet, A.B., et al., Studying the Endothelial Glycocalyx in vitro: What Is Missing? Front Cardiovasc Med, 2021. 8: p. 647086.

63. Tarbell, J.M. and L.M. Cancel, The glycocalyx and its significance in human medicine. J Intern Med, 2016. 280(1): p. 97–113.

64. Spillings, B.L., et al., Host glycocalyx captures HIV proximal to the cell surface via oligomannose-GlcNAc glycan-glycan interactions to support viral entry. Cell Rep, 2022. 38(5): p. 110296.

65. Hallak, L.K., et al., Glycosaminoglycan sulfation requirements for respiratory syncytial virus infection. J Virol, 2000. 74(22): p. 10508–13.

66. De Pasquale, V., et al., Heparan Sulfate Proteoglycans in Viral Infection and Treatment: A Special Focus on SARS-CoV-2. Int J Mol Sci, 2021. 22(12).

67. Cagno, V., et al., Heparan Sulfate Proteoglycans and Viral Attachment: True Receptors or Adaptation Bias? Viruses, 2019. 11(7).

68. Honigfort, D.J., et al., Glycocalyx crowding with mucin mimetics strengthens binding of soluble and virus-associated lectins to host cell glycan receptors. Proc Natl Acad Sci U S A, 2021. 118(40).

69. Weiss, R.J., J.D. Esko, and Y. Tor, Targeting heparin and heparan sulfate protein interactions. Org Biomol Chem, 2017. 15(27): p. 5656–5668.

70. Malhotra, R., et al., Isolation and characterisation of potential respiratory syncytial virus receptor(s) on epithelial cells. Microbes Infect, 2003. 5(2): p. 123–33.

71. Tayyari, F., et al., Identification of nucleolin as a cellular receptor for human respiratory syncytial virus. Nat Med, 2011. 17(9): p. 1132–5.

72. Xu, D. and J.D. Esko, Demystifying heparan sulfate-protein interactions. Annu Rev Biochem, 2014. 83: p. 129–57.

73. Cheng, F., et al., Nucleolin is a nuclear target of heparan sulfate derived from glypican-1. Exp Cell Res, 2017. 354(1): p. 31–39.

74. Bourgeois, C., et al., Heparin-like structures on respiratory syncytial virus are involved in its infectivity in vitro. J Virol, 1998. 72(9): p. 7221–7.

75. Karger, A., U. Schmidt, and U.J. Buchholz, Recombinant bovine respiratory syncytial virus with deletions of the G or SH genes: G and F proteins bind heparin. J Gen Virol, 2001. 82(Pt 3): p. 631–640.

76. Kim, M. and Y. Kim, NMR Structural Study of Syndecan-4 Transmembrane Domain with Cytoplasmic Region. Molecules, 2023. 28(23).

77. Manon-Jensen, T., Y. Itoh, and J.R. Couchman, Proteoglycans in health and disease: the multiple roles of syndecan shedding. FEBS J, 2010. 277(19): p. 3876–89.

78. Elfenbein, A. and M. Simons, Syndecan-4 signaling at a glance. J Cell Sci, 2013. 126(Pt 17): p. 3799–804.

79. Vittum, Z., S. Cocchiaro, and S.A. Mensah, Basal endothelial glycocalyx’s response to shear stress: a review of structure, function, and clinical implications. Front Cell Dev Biol, 2024. 12: p. 1371769.

80. Zhu, Z., et al., Syndecan-4 is the key proteoglycan involved in mediating sepsis-associated lung injury. Heliyon, 2023. 9(8): p. e18600.

