## supplementary data for "Evidence that the cell glycocalyx envelops respiratory syncytial virus (RSV) particles that form on the surface of RSV-infected human airway cells"

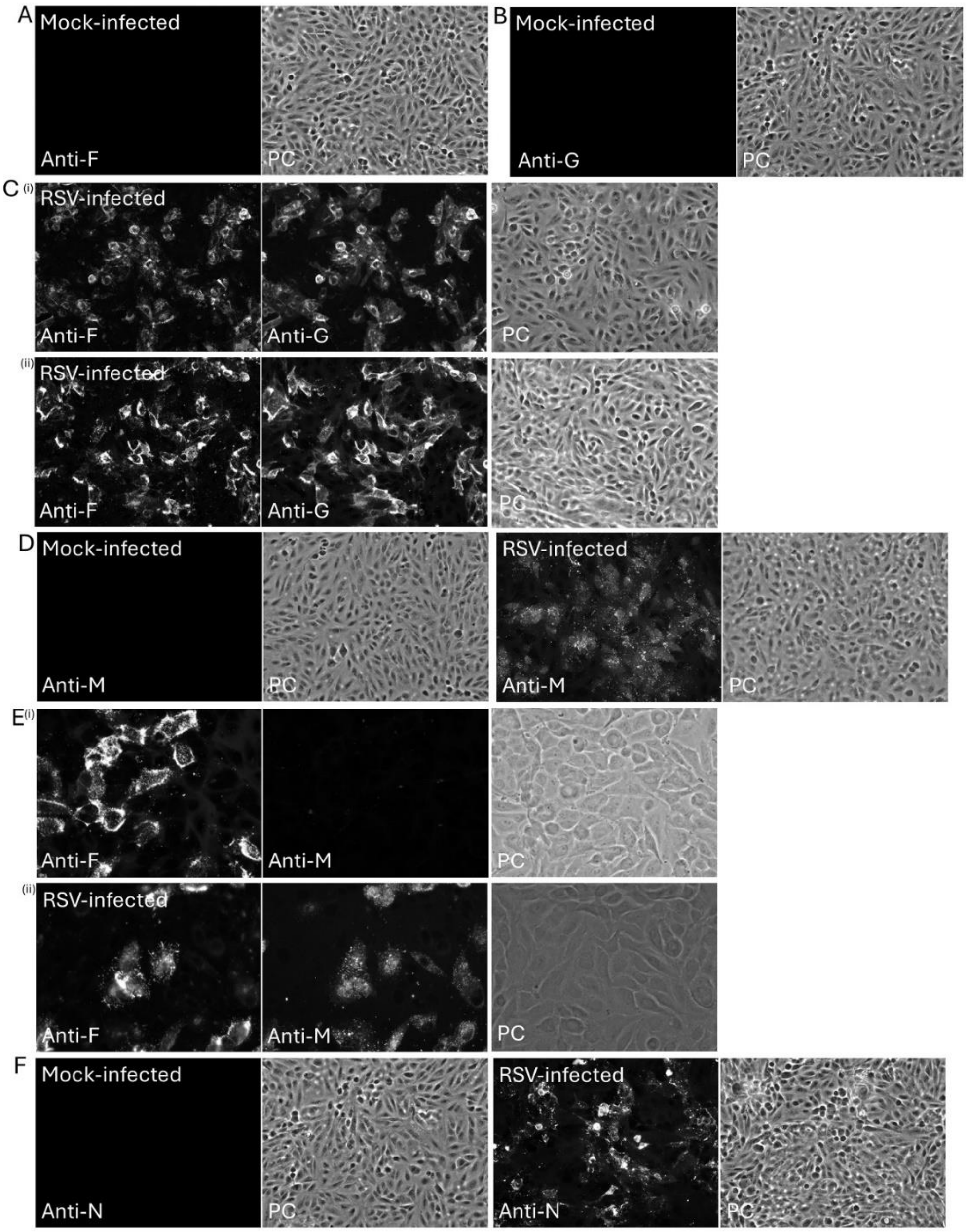


**SFigure 1. Distribution of RSV-infected cells at 24 hrs post infection.** A549 cell monolayers were either mock-infected or RSV-infected as indicated and the cells harvested at 24 hrs post-infection. **(A and B)** Mock-infected cells were permeabilised and stained with either anti-G or anti-F as indicated, and the RSV-infected cells were **(C)** (i) non-permeabilised and (ii) permeabilised and co-stained with anti-F and anti-G. The cells were imaged using immunofluorescence (IF) microscopy and phase contrast (PC) microscopy (objective x20 magnification). **(D)** Mock-infected and RSV-infected cells were permeabilised and stained with anti-M and the cells were imaged using IF and PC microscopy (objective x20 magnification). **(E)** RSV-infected cells were (i) non-permeabilised and (ii) permeabilised and co-stained with anti-M and anti-F and the cells were imaged using IF and PC microscopy (objective x40 magnification). **(F)** Mock-infected and the RSV-infected cells were stained with anti-N and the cells were imaged using IF and PC microscopy (objective x20 magnification).

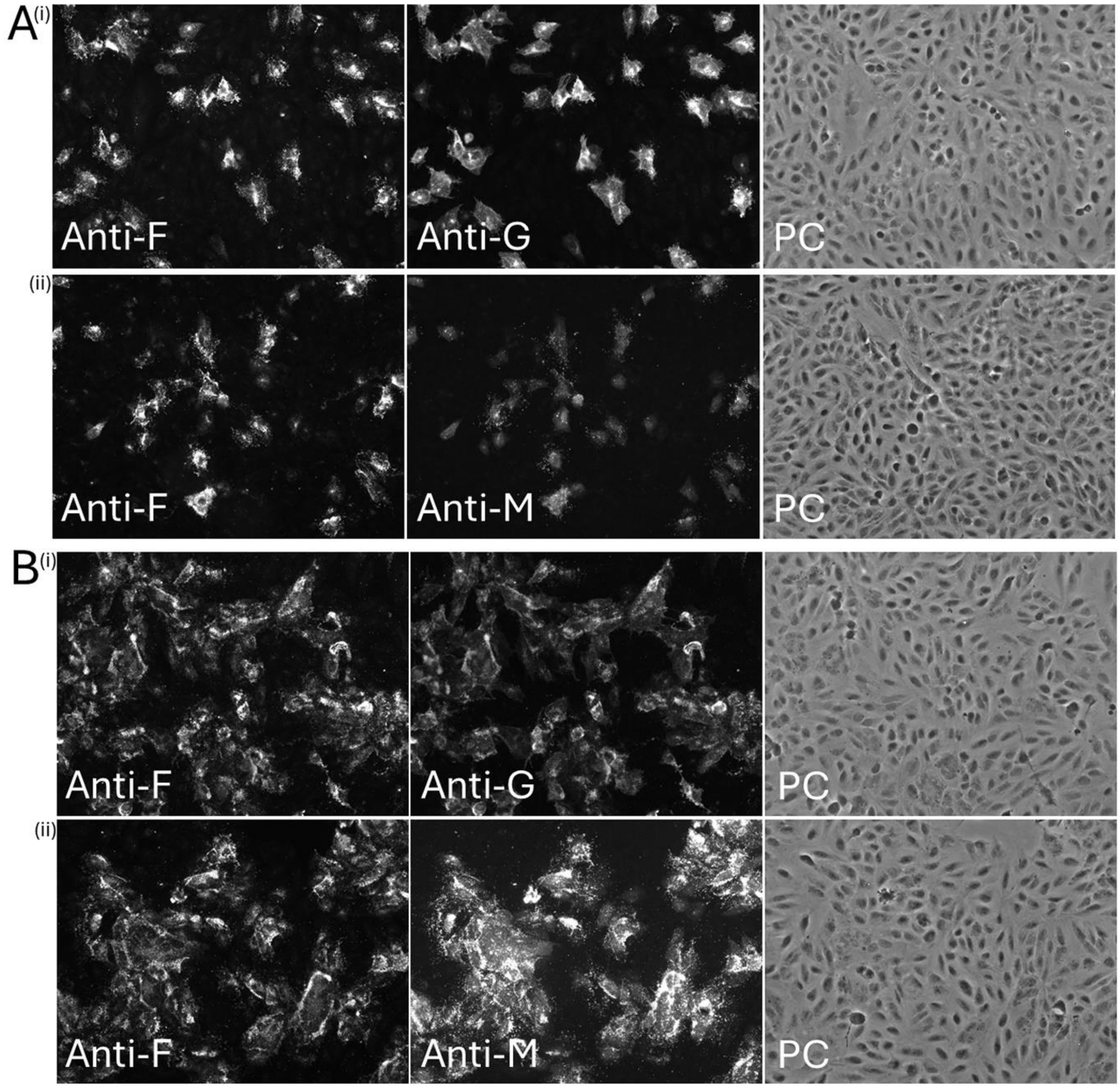


**SFigure 2. Distribution of RSV-infected cells at 24 and 42 hrs post-infection in the A549 cell monolayers.** A549 cells cell monolayers were (i) mock-infected and (ii) RSV-infected using a multiplicity of infection of 0.05 and at **(A)** 24 hrs post-infection and **(B)** 48 hrs post-infection cells were permeabilized and co-stained with (i) anti-F and anti-G and (ii) anti-F and anti-M and imaged using immunofluorescence microscopy and phase contrast microscopy (PC) (objective x20 magnification).

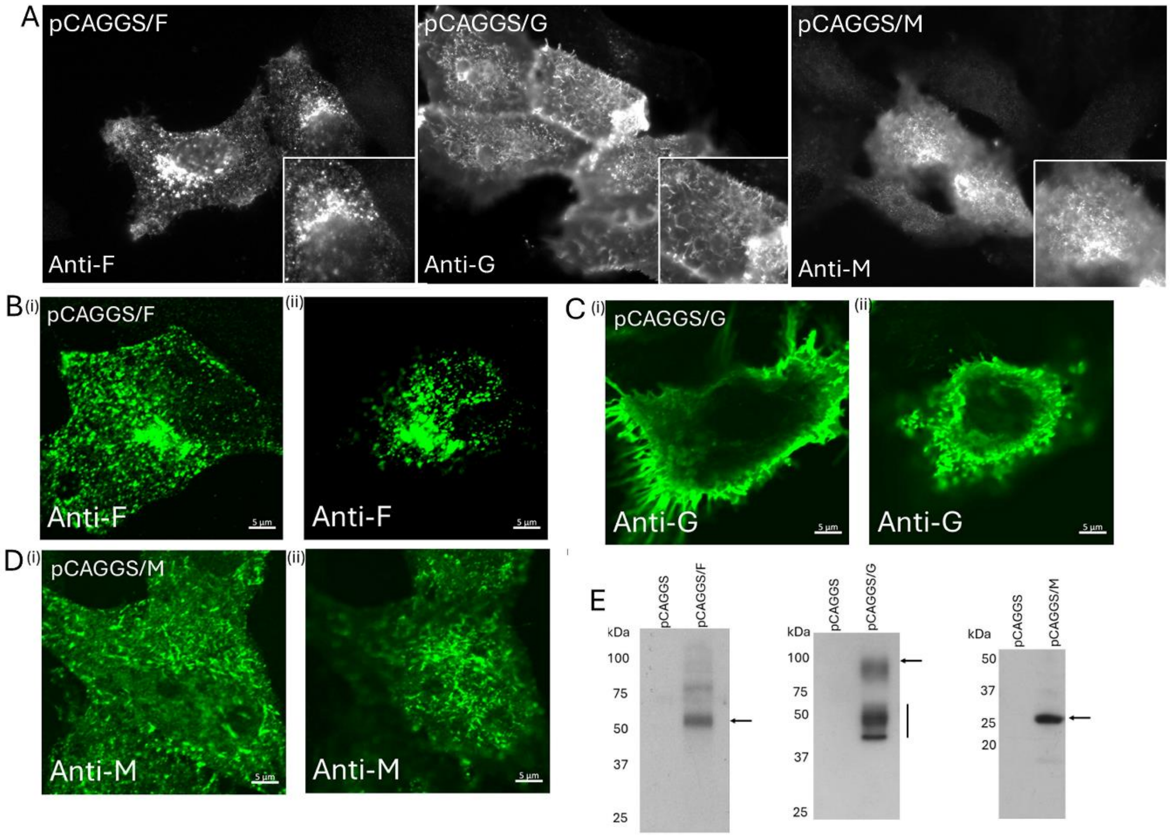


**SFigure 3. The respective staining pattern of the recombinant expressed G, F and M proteins in HEp2 cells.** HEp2 cells were transfected with pCAGGS/F and pCAGGS/G and the non-permeabilised and stained with anti-F and anti-G respectively, and the pCAGGS/M-transfected cells were permeabilised and stained with anti-M as indicated**.** **(A)** The stained cells were imaged using immunofluorescence microscopy (objective x40 magnification). Insets are enlarged images of representative antibody-stained cells. **(B-D)** cells were transfected with **(B)** pCAGGS/F and **(C)** pCAGGS/G and non-permeabilised and stained with anti-F and anti-G as indicated and **(D)** with pCAGGS/M and the cells were permeabilised and stained with anti-M as indicated. Representative cells were imaged using confocal microscopy at a focal plane that allowed imaging of the (i) the cell periphery and (ii) cell top. **(E)** Cells lysates were prepared from pCAGGS, pCAGGS/F, pCAGGS/G pCAGGS/M as indicated and immunoblotted with the appropriate antibody. Protein species of the expected sizes for the respective virus protein are indicated (black arrow). A smaller protein smeared band corresponding to the partially mature G protein is also indicated (black line). The cell lysates obtained from cells transfected with pCAGGS provide a negative control. In all panels the cells were examined at 18 hrs post-transfection.
